# Quorum-sensing *agr* system of *Staphylococcus aureus* primes gene expression for protection from lethal oxidative stress

**DOI:** 10.1101/2023.06.08.544038

**Authors:** Magdalena Podkowik, Andrew I. Perault, Gregory Putzel, Andrew Pountain, Jisun Kim, Ashley Dumont, Erin Zwack, Robert J. Ulrich, Theodora K. Karagounis, Chunyi Zhou, Andreas F. Haag, Julia Shenderovich, Gregory A. Wasserman, Junbeom Kwon, John Chen, Anthony R. Richardson, Jeffrey N. Weiser, Carla R. Nowosad, Desmond S. Lun, Dane Parker, Alejandro Pironti, Xilin Zhao, Karl Drlica, Itai Yanai, Victor J. Torres, Bo Shopsin

## Abstract

The *agr* quorum-sensing system links *Staphylococcus aureus* metabolism to virulence, in part by increasing bacterial survival during exposure to lethal concentrations of H_2_O_2_, a crucial host defense against *S. aureus*. We now report that protection by *agr* surprisingly extends beyond post-exponential growth to the exit from stationary phase when the *agr* system is no longer turned on. Thus, *agr* can be considered a constitutive protective factor. Deletion of *agr* increased both respiration and fermentation but decreased ATP levels and growth, suggesting that Δ*agr* cells assume a hyperactive metabolic state in response to reduced metabolic efficiency. As expected from increased respiratory gene expression, reactive oxygen species (ROS) accumulated more in the *agr* mutant than in wild-type cells, thereby explaining elevated susceptibility of Δ*agr* strains to lethal H_2_O_2_ doses. Increased survival of wild-type *agr* cells during H_2_O_2_ exposure required *sodA*, which detoxifies superoxide. Additionally, pretreatment of *S. aureus* with respiration-reducing menadione protected Δ*agr* cells from killing by H_2_O_2_. Thus, genetic deletion and pharmacologic experiments indicate that *agr* helps control endogenous ROS, thereby providing resilience against exogenous ROS. The long-lived “memory” of *agr*-mediated protection, which is uncoupled from *agr* activation kinetics, increased hematogenous dissemination to certain tissues during sepsis in ROS-producing, wild-type mice but not ROS-deficient (Nox2^−/−^) mice. These results demonstrate the importance of protection that anticipates impending ROS-mediated immune attack. The ubiquity of quorum sensing suggests that it protects many bacterial species from oxidative damage.

## Introduction

Innate, bactericidal immune defenses and antimicrobials act, at least in part, by stimulating the accumulation of reactive oxygen species (ROS) in bacteria (1, 2). Thus, understanding how *Staphylococcus aureus* and other bacterial pathogens manage ROS-mediated stress has important implications for controlling infections.

Knowledge of factors that govern the biology of ROS has advanced considerably in recent years. For example, studies have centered on how specific metabolic features, such as aerobic respiration, affect killing by ROS (3, 4), and small-molecule enhancers of ROS-mediated lethality are emerging (5, 6). Less well characterized is how defense against ROS and metabolism changes integrate with the virulence regulatory network that promotes *S. aureus* pathogenesis. The *agr* quorum-sensing system provides a way to study this dynamic: *agr* is a major virulence regulator that responds to oxidative stress (H_2_O_2_). The response occurs through a redox sensor in AgrA that attenuates *agr* activity, thereby increasing expression of glutathione peroxidase (BsaA), an enzyme that detoxifies ROS (7). Whether protection from ROS also occurs from positive *agr* action is unknown and likely to be an important issue in development of Agr-targeted therapies (8).

In cultured *S. aureus*, *agr* governs the expression of ∼200 genes. Its two-part regulatory role is characterized by 1) increased post-exponential-phase production of toxins and exoenzymes that facilitate dissemination of bacteria via tissue invasion, and 2) decreased production of cell surface and other proteins that facilitate adherence, attachment, biofilm production, and evasion of host defenses (9, 10). Thus, *agr* coordinates a switch from an adherent state to an invasive state at elevated bacterial population density. The invasive state would be facilitated by protection from host defense.

The *agr* locus consists of two divergent transcription units driven by promoters P2 and P3 (11). The P2 operon encodes the quorum-signaling module, which contains four genes, *agrB, agrD, agrC*, and *agrA*. AgrC is a receptor histidine kinase, and AgrA is a DNA-binding response regulator. AgrD is an autoinducing, secreted peptide derived from a pro-peptide processed by AgrB. The autoinducing peptide binds to and causes autophosphorylation of the AgrC histidine kinase, which phosphorylates and activates the DNA-binding AgrA response regulator. AgrA then stimulates transcription from the P2 (RNAII) and P3 (RNAIII) promoters. RNAIII is a regulatory RNA that additionally contains the gene for delta-hemolysin (*hld*). The DNA-binding domain of AgrA contains an intramolecular disulfide switch (7). Oxidation leads to dissociation of AgrA from DNA, thereby preventing an AgrA-mediated down-regulation of the BsaA peroxidase.

When we used antimicrobials to study bacterial responses to lethal stress involving the accumulation of ROS, we found that inactivation (deletion) of *agr* reduces lethality arising from treatment with antimicrobials, such as fluoroquinolones, in a largely *bsaA*-dependent manner (12). Thus, oxidation sensing appears to be an intrinsic checkpoint that ameliorates the endogenous oxidative burden generated by certain antimicrobials. Surprisingly, deletion of *agr increases* the lethal effects of exogenous H_2_O_2_ (12), in contrast to the expected expression of the protective *bsaA* system (7). Thus, *agr* must help protect *S. aureus* from exogenous ROS, a principal host defense, through mechanisms other than *bsaA*.

In the present work we found that protection by wild-type *agr* against lethal concentrations of H_2_O_2_ was unexpectedly long-lived and 1) associated with decreased expression of respiration genes, and 2) potentially aided by defense systems that suppress the oxidative surge triggered by subsequent, high-level H_2_O_2_ exposure. The redox switch in AgrA, plus these additional protective properties, indicate that *agr* increases resilience to oxidative stress in *S. aureus* both when it is present and when it is absent. Thus, *agr* integrates protection from host defense into the regulation of staphylococcal virulence.

## Results

### agr protects S. aureus from lethal concentrations of H2O2 throughout the growth cycle

Because *agr* is a quorum-sensing regulon, maximal *agr* activity occurs during exponential growth (Figure 1. figure supplement 1) and is followed by a sharp drop during stationary phase (12, 13). Surprisingly, protection from H_2_O_2_ toxicity by wild-type *agr*, assessed by comparison with an *agr* deletion mutant, was observed throughout the growth cycle (Figure 1A). Indeed, maximal protection occurred shortly after overnight growth, long after induction and expression of *agr* transcripts. Comparison of survival rates of *Δagr* mutant and wild-type cells, following dilution of overnight cultures and regrowth for 1 h prior to challenge with 20 mM H_2_O_2_, revealed an initial rate of killing that was ∼1,000-fold faster for the *Δagr* mutant (Figure 1B). Peroxide concentration dependence was observed up to 10 mM during a 60-min treatment; at that point, mutant survival was about 100-fold lower (Figure 1C). Complementation tests confirmed that the *agr* deletion elevated killing by H_2_O_2_ (Figure 1D).

**Figure 1.**
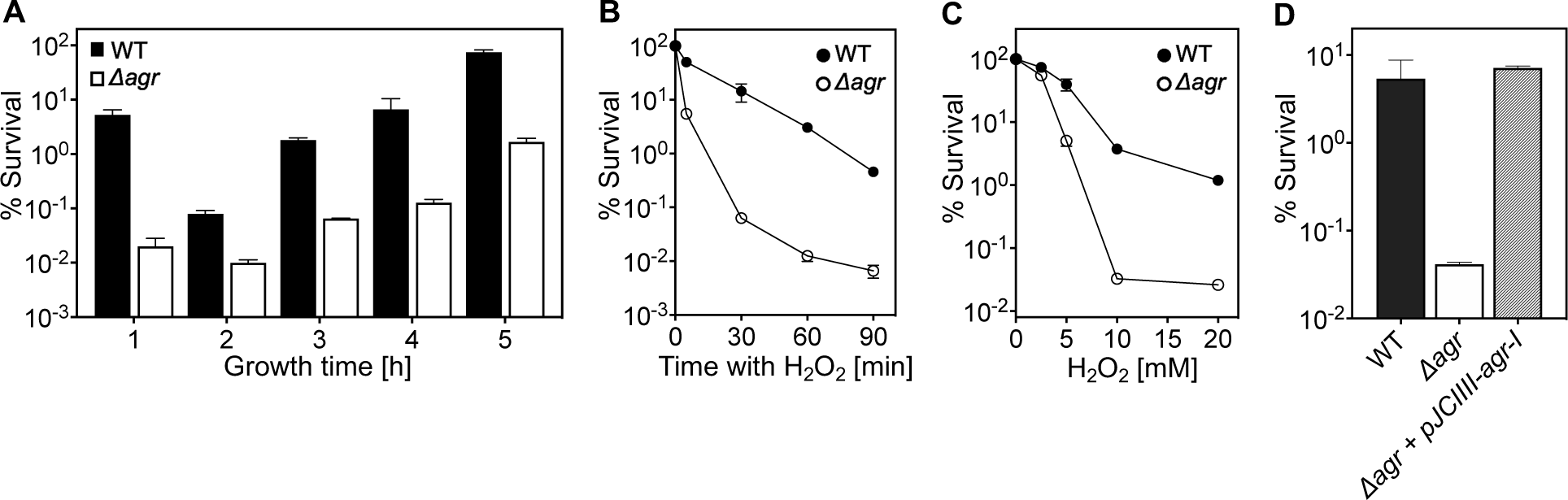
*agr* protects from killing by H_2_O_2_ throughout the growth cycle. (A) Effect of culture growth phase. Overnight cultures of *S. aureus* LAC wild-type (WT, BS819) or *Δagr* (BS1348) were diluted (OD_600_∼0.05) into fresh TSB medium and grown with shaking from early exponential (1 h, OD_600_∼0.15) through late log (5 h, OD_600_∼4) phase. At the indicated times, early (undiluted) and late exponential phase cultures (diluted into fresh TSB medium to OD_600_∼0.15) were treated with H_2_O_2_ (20 mM). After 60 min, aliquots were removed, serially diluted, and plated for determination of viable counts. Percent survival was calculated relative to a sample taken at the time of H_2_O_2_ addition. (B) Kinetics of killing by H_2_O_2_. Wild-type and *Δagr* mutant strains were grown to early exponential (OD_600_∼0.15) and treated with 20 mM H_2_O_2_ for the times indicated, and percent survival was determined by plating. (C) Effect of H_2_O_2_ concentration on survival. Cultures prepared as in panel B were treated with the indicated peroxide concentrations for 60 min prior to plating and determination of percent survival. (D) Complementation of *agr* deletion mutation. Cultures of wild-type (WT) cells (BS819), Δ*agr* mutant (BS1348), and complemented Δ*agr* mutant carrying a chromosomally integrated wild-type operon (pJC1111-*agrI*) were treated with 20 mM H_2_O_2_ for 60 min followed by plating to determine percent survival. Data represent the means ± S.D. from biological replicates (*n* = 3).

We also monitored the time required for the wild-type *agr* survival advantage against H_2_O_2_ to manifest itself (Figure 1. figure supplement 2). Overnight cultures were not readily killed by H_2_O_2_, as expected from previous results with other lethal stressors (14). Following dilution to fresh medium, wild-type survival dropped gradually, while mutant survival, although lower, was constant for 20 min. By 40 min, mutant survival exhibited a precipitous 10-fold drop not seen with wild-type cells (Figure 1. figure supplement 2). This drop in mutant survival correlated temporally with changes in cell density (Figure 1. figure supplement 2); i.e., the first cell division following dilution to fresh medium. Overall, the *agr*-mediated survival advantage during H_2_O_2_ exposure was absent in stationary-phase cells and small during lag phase (before exponential growth resumes), but it increased markedly during early growth.

Lag-time differences between strains were more obvious in experiments using less complex, chemically defined medium (CDM) with highly diluted starting cultures and automated growth analysis (Figure 1. figure supplement 3). In CDM, wild-type cells divided within ∼150 min, while the lag times with the *Δagr* mutant were more than 205 min (in Tryptic Soy Broth the lag time is 30 min for both). These observations suggest a novel *agr*-mediated decrease in time to enter exponential growth following dilution of stationary phase cultures. The poor killing of *agr* mutant cells by H_2_O_2_ early in lag phase is consistent with other work in which cells experiencing long lag times are less readily killed (15), presumably due to remaining longer in a dormant, protected state. To focus on effects during growth, subsequent experiments were performed after incubation of overnight cultures for 1 h in fresh Tryptic Soy Broth unless otherwise specified.

The elevating effect of *agr* inactivation on H_2_O_2_-mediated lethality was observed across a variety of *S. aureus* strains, although differences in wild-type survival were observed (Figure 1. figure supplement 4). Thus, *agr*-mediated protection from H_2_O_2_ appears to be common among *S. aureus* lineages.

### Expression of RNAIII and repression of Rot is required for protection from H2O2-mediated lethality

Δ*rnaIII* and Δ*agr* mutants showed identical loss of protection from H_2_O_2_-mediated killing (Figure 2A), indicating that protection is RNAIII-dependent. Since RNAIII represses translation of the downstream regulator Rot (16), a transcription factor having a key role in *agr* regulation of staphylococcal virulence, we also examined the effects of *rot* on the protective action of *agr* against H_2_O_2_. When the wild-type strain, a Δ*agr* mutant, a Δ*rot* mutant, and a Δ*agr* Δ*rot* double mutant were compared for survival following treatment with 20 mM H_2_O_2_, survival of the Δ*agr* Δ*rot* double mutant phenocopied that of the wild-type strain (Figure 2B): the *rot* deletion reversed the effect of an *agr* deficiency. These data are consistent with *agr* activity allowing induction of *rot*-repressed genes important for protection from peroxide (RNAIII repression of the Rot repressor).

**Figure 2.**
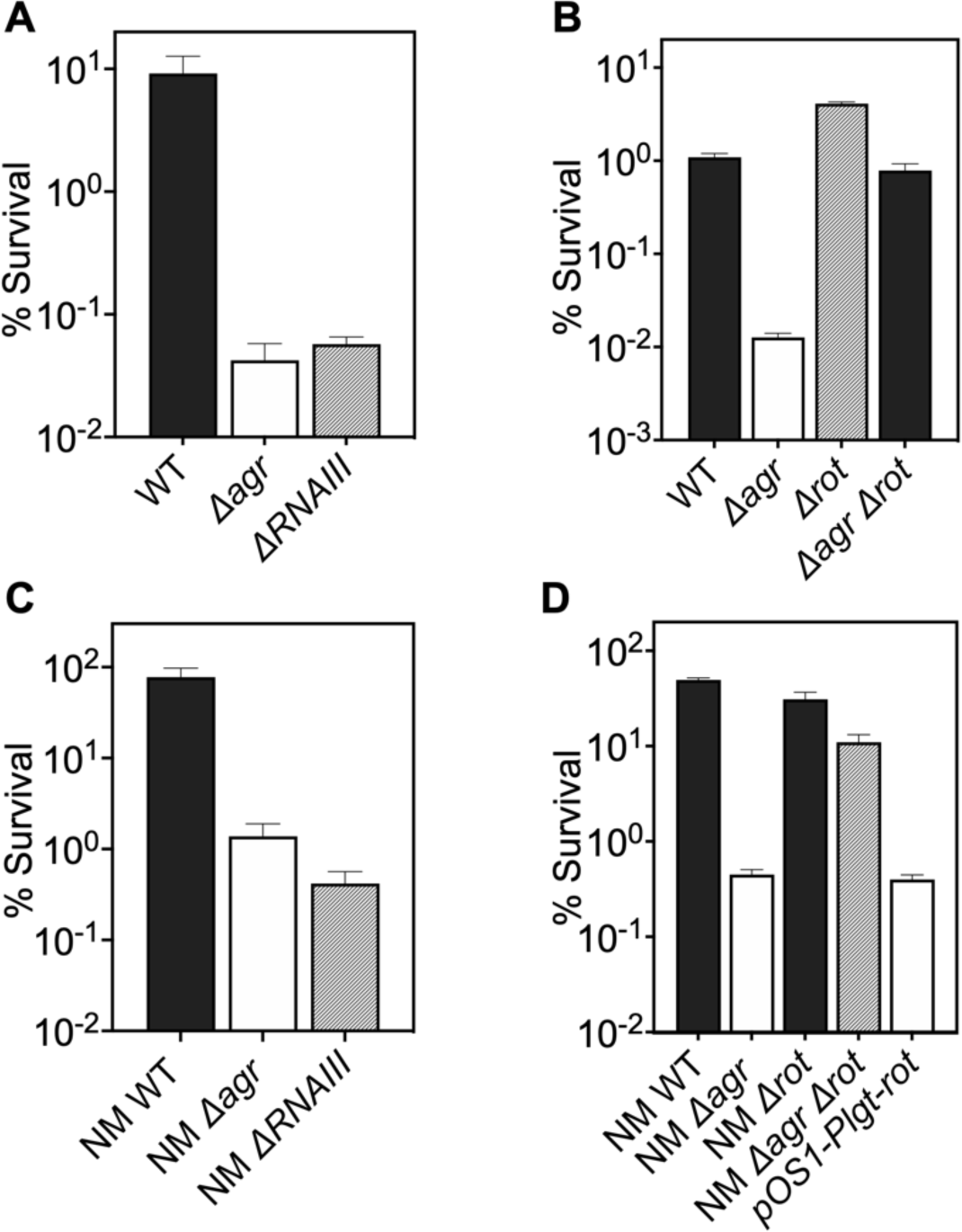
Involvement of *RNAIII* and *rot-*dependent pathways in *agr-*mediated protection from H_2_O_2_-mediated killing. Cultures were grown for 1 h following dilution from overnight cultures to early log phase (OD_600_∼0.15) and then treated with 20 mM H_2_O_2_ for 60 min before determination of percent survival by plating and enumeration of colonies. (A) Wild-type LAC (WT, BS819), *Δagr* mutant (BS1348), and *ΔrnaIII* mutant (GAW183). (B) *Δrot* and *Δagr Δrot* double mutant (BS1302). (C) Wild-type (WT) strain Newman (NM, BS12), *Δagr* mutant (BS13), and *ΔRNAIII* mutant (BS669). (D) Overexpression of *rot*. Rot was expressed from a plasmid-borne wild-type *rot* (pOS1-*Plgt-rot*, strain VJT14.28). Data represent the means ± S.D. from biological replicates (*n* = 3).

When a low-copy-number plasmid expressing *rot* was introduced into a wild-type strain, the transformant was more readily killed by H_2_O_2_, indicating that expression of *rot* is sufficient for increased lethality (Figs. 2C-D). These data suggest that wild-type Rot down-regulates expression of protective genes. The observed epistatic effect of *agr* and *rot* did not apply to other downstream, potentially epistatic regulators, such as *saeRS*, *mgrA*, and *sigB* (Figure 2—figure supplement 1) (17). Thus, the epistatic relationship between *agr* and protection from H_2_O_2_ appears to be *rot*-specific.

### agr-mediated protection from H_2_O_2_ stress is kinetically uncoupled from agr activation

Since *agr*-mediated protection from H_2_O_2_ occurs throughout the growth cycle, it was possible that protection arises from constitutive, low-level *agr* expression rather than from autoinduction and thereby quorum sensing. To test for a requirement of quorum in the *agr*-mediated oxidative-stress phenotypes, we characterized the role of *agr* activation using a mixed culture strategy in which one strain, an in-frame deletion mutant of *agrBD*, is activated *in trans* by AIP produced by a second, Δ*rnaIII* mutant strain (Figure 3A). The AIP-responsive Δ*agrBD* strain carried an intact RNAIII, while the Δ*rnaIII* mutant was wild-type for *agrBD*. As shown in Figure 3B, hemolytic activity (a marker for RNAIII) of the Δ*agrBD* mutant was restored by mixing it with the Δ*rnaIII* mutant strain that secreted AIP into the surrounding medium. This result confirmed that *agrCA*-directed *trans*-activation of RNAIII by AIP remained intact in the Δ*agrBD* mutant.

**Figure 3.**
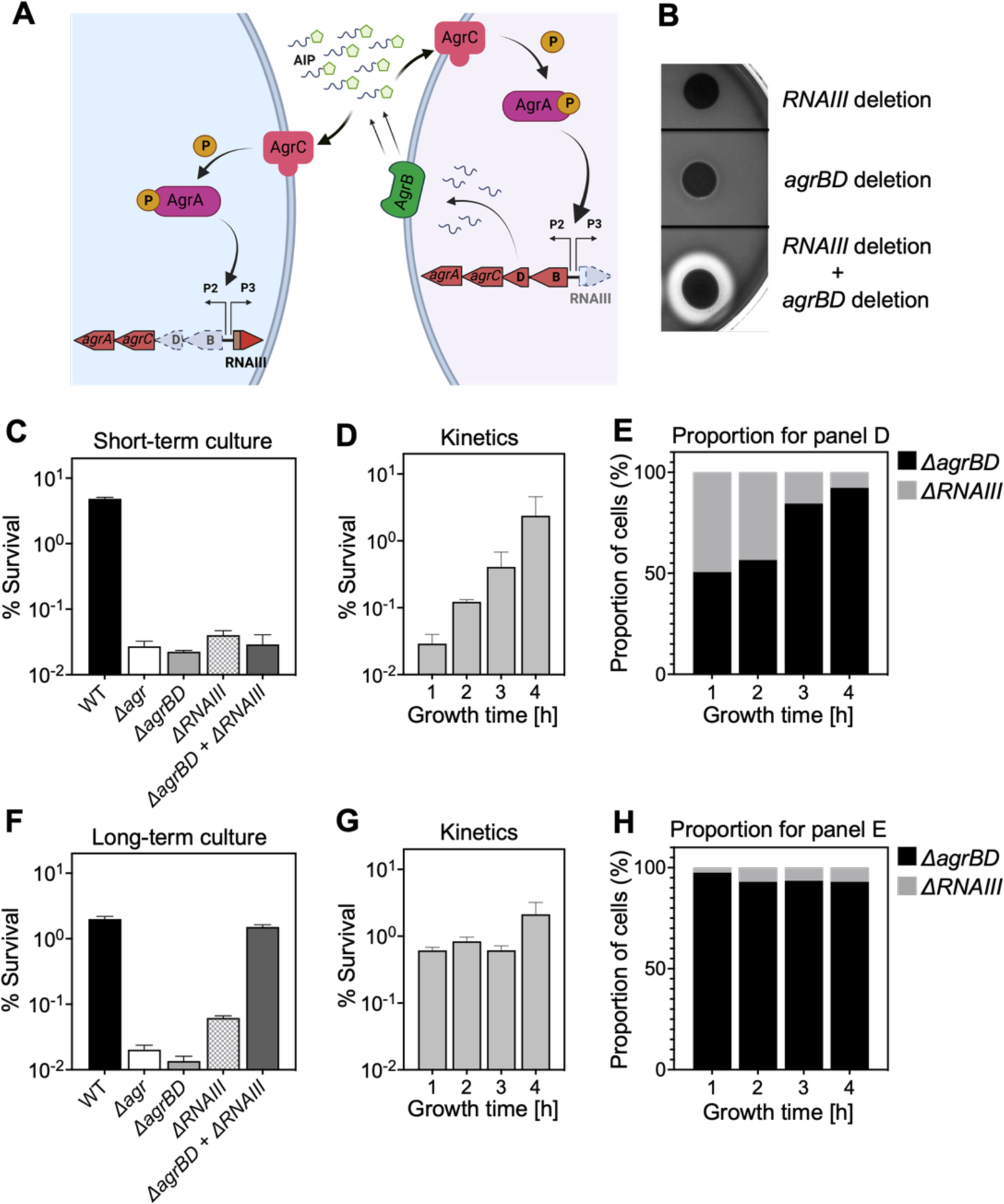
*agr*-mediated protection from H_2_O_2_ stress is uncoupled from *agr* activation kinetics. (A) Assay design. An Δ*agrBD* deletion mutant (GAW130) was complemented *in trans* by the autoinducing product (AIP) of AgrBD in an *ΔrnaIII* (GAW183) mutant that produces AIP endogenously; AgrC activation in the Δ*agrBD* strain leads to downstream activation of RNAIII. *The agrBD* strain, engineered in-frame to avoid polar effects on downstream genes *agrC* and *agrA*, senses but does not produce autoinducer. The Δ*rnaIII* mutant, constructed by replacement of *rnaIII* with a cadmium resistance cassette (*rnaIII*::*cadA*), produces autoinducer but lacks RNAIII, the effector molecule of *agr*-mediated phenotypes with respect to H_2_O_2_. The image was created using BioRender (BioRender.com). (B) Trans-activation demonstrated by hemolysin activity on sheep blood agar plates. Bottom of figure shows zone of clearing (hemolysin activity) after mixing 10^8^ Δ*agrBD* CFU with an equal number of Δ*rnaIII.* Zone of clearance is a consequence of *AgrC* receptor activation *in trans* by AIP produced by the Δ*rnaIII* mutant. (C) Absence of trans-activation with short-term culture. The wild-type strain RN6734 (WT, BS435), Δ*rnaIII* (GAW183), Δ*agrBD* (GAW130), and Δ*rnaIII* and Δ*agrBD* mutants were mixed 1:1 immediately before growth from overnight culture. Overnight cultures were diluted (OD_600_∼0.05) into fresh TSB medium, mixed, and grown to early log phase (OD_600_∼0.15) when they were treated with 20 mM H_2_O_2_ for 60 min and assayed for percent survival by plating. (D) Kinetics of killing by H_2_O_2._ Survival assays employing *ΔrnaIII* and *ΔagrBD* mixtures, performed as in panel C, but grown from early exponential (1 h, OD_600_∼0.15) through late log (5 h, OD_600_∼4) phase in TSB. Cultures were treated with H_2_O_2_ (20 mM for 1 h) at the indicated time points. (E) Proportion of mixed population for panel D represented by each mutant after incubation. The Δ*agrBD* mutant contained an erythromycin-resistance marker to distinguish the strains following plating of serial dilutions on TS agar with or without erythromycin (5 μg/ml). Data represent the means ± S.D. from biological replicates (*n* = 3). (F) Trans-activation during long-term culture. The wild-type strain RN6734 (WT, BS435), Δ*rnaIII* (strain GAW183), Δ*agrBD* (strain GAW130), and Δ*rnaIII* and Δ*agrBD* mutants mixed 1:1 prior to overnight culture. Survival assays employing Δ*rnaIII* and Δ*agrBD* mixtures, performed as in panel C. (G) Kinetics of killing by H_2_O_2_. Survival assays employing Δ*rnaIII* and Δ*agrBD* mixtures, performed as in panel D. Cultures were treated with H_2_O_2_ (20 mM for 1 h) at the indicated time points. (H) Proportion of mixed population for panel G represented by each mutant after incubation, performed as in panel E. Data represent the means ± S.D. from biological replicates (*n* = 3).

Mixed culture tests using these mutants, scored by differential plating for the presence of an erythromycin resistance marker in the Δ*agrBD* mutant, showed no protection from lethality of H_2_O_2_ when the two strains were mixed 1:1 immediately prior to growth from stationary phase (Figure 3C). Autoinducer accumulated during subsequent growth, activating *agr* expression and commencing protection from exogenous H_2_O_2_ (Figs. 3D-E). During H_2_O_2_ treatment, the percentage of the Δ*agrBD* mutant (*rnaIII*^+^) increased while the percentage of the Δ*rnaIII* mutant decreased; this cis-acting result is consistent with the idea that pathways downstream from RNAIII, such as those regulated by *rot*, are the primary drivers of *agr*-mediated protection from H_2_O_2_. These results confirm an intimate link between *agr*-mediated protection and the quorum-controlled *agr* gene expression program of late exponential phase. However, after an overnight co-culture of the Δ*rnaIII* and Δ*agrBD* mutant strains, the Δ*agrBD* mutant demonstrated the same degree of protection expected for wild-type cells during exposure to H_2_O_2_ (Figure 3F-H). Thus, protection by *agr* after overnight co-culture extends to growth resumption from stationary phase, prior to reaching quorum, and therefore protection is uncoupled from the constraint of strict cell-density dependence. These results indicate that protection lasts long after maximal transcription of *agr*, when *agr* expression has largely halted (12, 13). This phenomenon is a critical feature of the *agr* system not appreciated in previous analyses of *agr* activation kinetics.

### agr deficiency increases transcription of genes involved in respiration and overflow metabolism in the absence of stress

To explore mechanisms underlying protection from H_2_O_2_, we performed RNA-seq with the Δ*agr* and wild-type strains after growth to late exponential growth phase, a point when *agr* expression is maximal. As expected, *agr* up-regulated the transcription of many known virulence genes (Supplementary file 1). The Δ*agr* strain showed elevated expression of genes involved in respiration (*cydA*, *qoxA-D*) and fermentation (18, 19), including *nrdGD,* alcohol dehydrogenases (*adhE* and *adh1*) and lactate dehydrogenases (*ldh*, *ddh*) (Figure 4A and Supplementary file 1). Increased respiration and fermentation are expected to increase energy generation. However, metabolic modeling of transcriptomic data showed a ∼30% reduction in tricarboxylic acid (TCA) cycle and lactate flux per unit of glucose taken up by the Δ*agr* mutant (Figure 4B, Supplementary file 2). Additionally, intracellular ATP levels were ∼50% lower in the *Δagr* mutant compared to the wild-type control, suggesting reduced metabolic efficiency during exponential growth (Figure 5A). Moreover, although the *agr* deletion has little effect on growth in the rich medium in which RNA-seq was performed (20), analysis in nutrient-constrained medium (CDM) revealed decreased growth rate and yield of the Δ*agr* mutant relative to wild-type *S. aureus* (Figure 1—figure supplement 3). Collectively, these data suggest that Δ*agr* increases respiration and fermentation to compensate for low metabolic efficiency. Consistent with this idea, *agr* deficiency also increases ATP-yielding carbon “overflow” pathways, as evidenced by increased acetate production (Figure 5B) (21, 22). The increase in accumulated acetate in the culture medium during exponential growth was largely consumed after 24 h of growth (Figure 5B). Thus, Δ*agr* mutants exhibit TCA cycle proficiency (20) and, despite some expense of efficiency, an increased catabolism of acetate.

**Figure 4.**
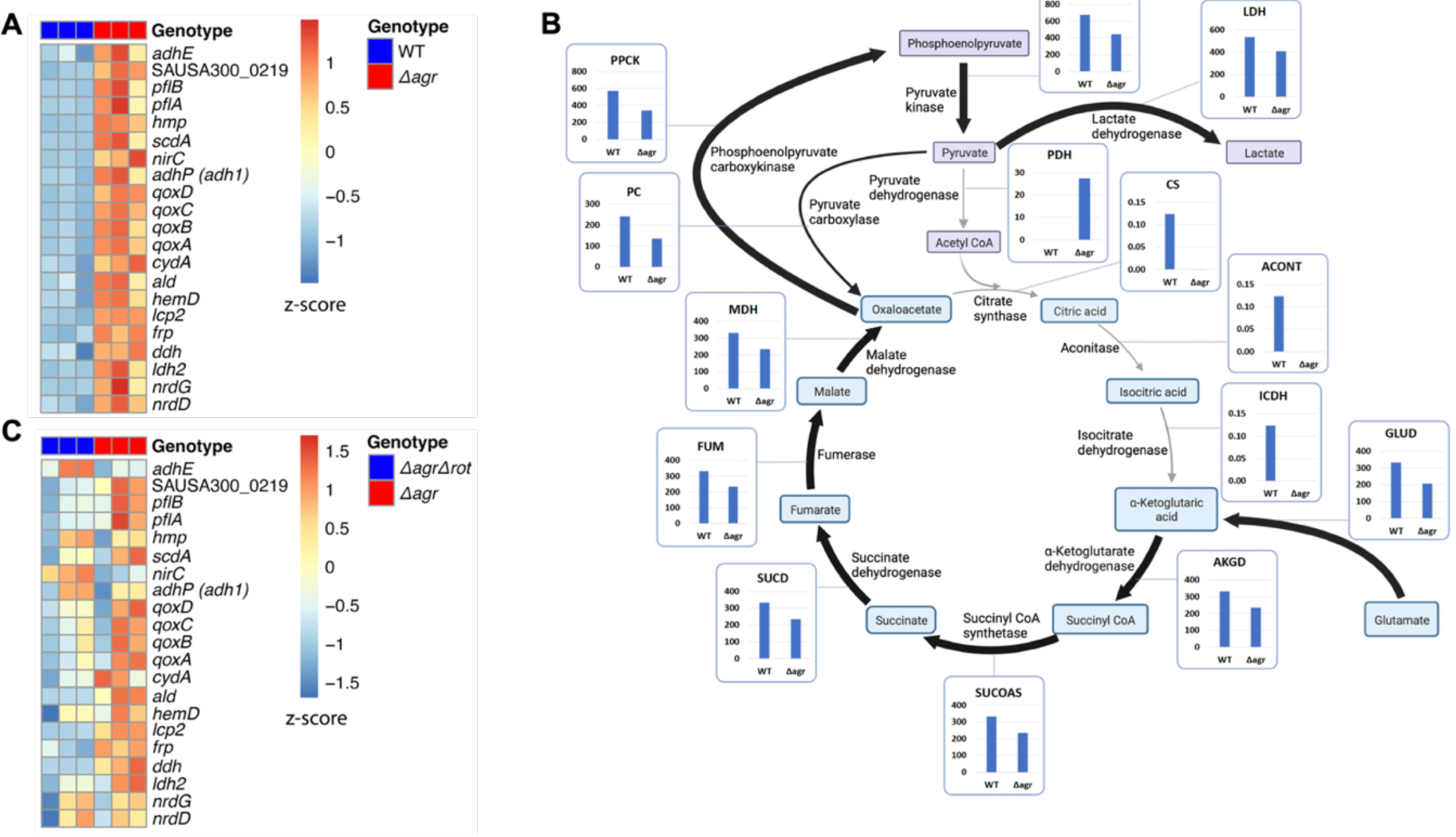
Association of *agr* deficiency with increased expression of respiration and fermentation genes during aerobic growth. (A) Relative expression of respiration and fermentation genes. RNA-seq comparison of *S. aureus* LAC wild-type (WT, BS819) and Δ*agr* mutant (BS1348) grown to late exponential phase (OD_600_∼4.0). Shown are significantly up-regulated genes in the Δ*agr* mutant (normalized expression values are at least twofold higher than in the wild-type). Heatmap colors indicate expression z-scores. RNA-seq data are from three independent cultures. See Supplementary file 1 for supporting information. (B) Schematic representation of *agr*-induced changes in metabolic flux, inferred from transcriptomic data (Supplementary file 1) by SPOT (Simplified Pearson correlation with Transcriptomic data). Metabolic intermediates and enzymes involved in catalyzing reactions are shown. The magnitude of the flux (units per 100 units of glucose uptake flux) is denoted by arrowhead thickness. Boxed charts indicate relative flux activity levels in wild-type versus Δ*agr* strains. Enzyme names are linked to abbreviations in boxed charts (e.g., lactate dehydrogenase, LDH). See Supplementary file 2 for supporting information. (C) RNA-seq comparison of an Δ*agr* Δ*rot* double mutant (BS1302) with its parental Δ*agr* strain (BS1348). Heatmap colors indicate expression z-scores. Sample preparation and figure labelling as for Figure 4A. See Supplementary file 3 for supporting information.

**Fig. 5.**
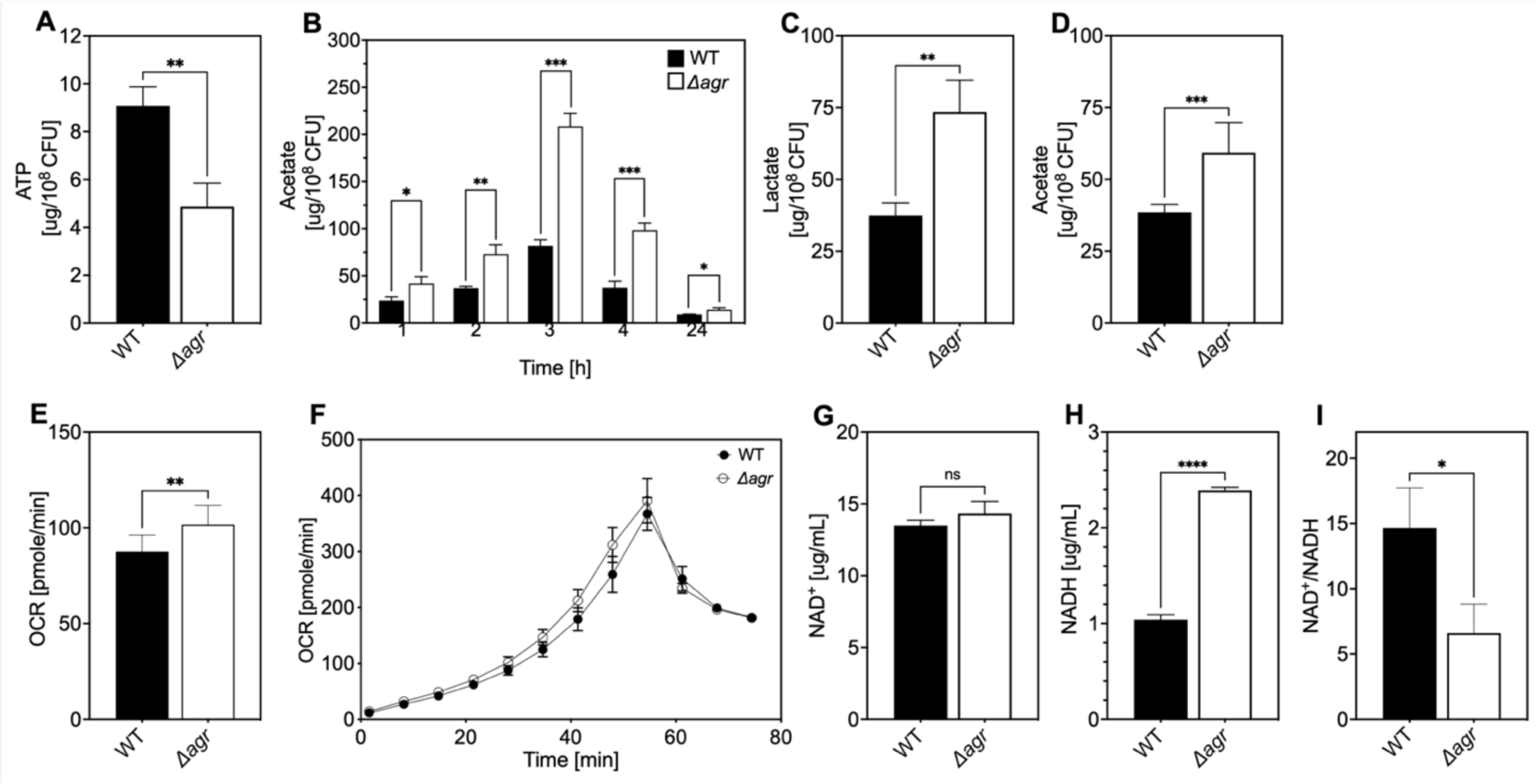
Association of *agr* deficiency with a metabolic flux shift toward fermentive metabolism during aerobic growth. (A) Intracellular ATP levels. Comparison of *S. aureus* LAC wild-type (WT, BS819) and Δ*agr* mutant (BS1348) strains for ATP expressed as µg/10^8^ cells after growth of cultures in TSB medium to late-exponential phase (OD_600_∼4.0). (B) Extracellular acetate levels. Samples were taken after 1, 2, 3, 4, and 24 h of growth in TSB medium; strains were wild-type (WT, BS819) and Δ*agr* mutant (BS1348). (C-D) Extracellular lactate and acetate levels during low oxygen culture. *S. aureus* LAC wild-type (WT, BS819) and Δ*agr* mutant (BS1348) were grown in TSB medium with suboptimal aeration to late-exponential phase (4 h, OD_600_∼4.0). (E-F) Oxygen consumption. Strains LAC wild-type (WT, BS819) and Δ*agr* mutant (BS1348) were compared using Seahorse XFp analyzer (F), and the rate of oxygen consumption (E) was determined from the linear portion of the consumption curve. Representative experiments from at least 3 independent assays are shown. (G-H) NAD^+^ and NADH levels. Colorimetric assay of NAD^+^ (G) and NADH levels (H) for *S. aureus* wild-type (WT, BS819) and Δ*agr* mutant (BS1348) after growth of cultures to late-exponential phase (OD_600_∼4.0). (I) NAD^+^/NADH ratio. For all panels, data points are the mean value ± SD (*n* = 3). \**P* < 0.05; \*\*\*\**P* < 0.0001, by Student’s two-tailed *t* test. Seahorse statistical significances are compared to TSB medium.

Differential transcription of selected genes was confirmed by RT-qPCR measurements (Figure 4—figure supplement 1) We also confirmed that respiration levels were lower (15%) in wild-type compared to Δ*agr* (Figure 5E-F). Although the stimulatory effect of the *agr* deletion on production of the fermentation product lactate was not observed in optimally aerated broth cultures after growth to late exponential growth phase, it was confirmed for organisms grown in broth under more metabolically demanding suboptimal aeration conditions (limitations in the rate of respiration when oxygen is limiting are expected to increase overall levels of fermentation) (Figure 5C). Overall, these results are consistent with transcription-level up-regulation of respiratory and fermentative pathways in *agr*-deficient strains.

Since respiration and fermentation generally increase NAD^+^/NADH ratios and since these activities are increased in *Δagr* strains (Figure 5C and E-F), we expected a higher NAD^+^/NADH ratio relative to wild-type cells. However, we observed a decrease in the NAD^+^/NADH ratio due to an increase in NADH accompanied by relative stability in NAD^+^ compared to wild-type. Collectively, these observations suggest that a surge in NADH accumulation and reductive stress in the Δ*agr* strain induces a burst in respiration, but levels of NADH are saturating, thereby driving fermentation under microaerobic condi6ons.

To help determine the metabolic fate of glucose, we measured glucose consumption and intracellular levels of pyruvate and TCA-cycle metabolites fumarate and citrate in the wild-type and *Δagr* mutant strains. At 4 h of growth to late-exponential phase, intracellular pyruvate and acetyl-CoA levels were increased in the *Δagr* mutant compared to wild-type strain, but levels of fumarate and citrate were similar (Figure 5—figure supplement 1D-E). Glucose was depleted after 4 h of growth, but glucose consumption after 3 h of growth (exponential phase) was increased in the *Δagr* mutant compared to the wild-type strain (Figure 5—figure supplement 1A). These observations, together with the decrease in the NAD^+^/NADH ratio and increase in acetate and lactate production described above, are consistent with a model in which respiration in Δ*agr* mutants is inadequate for 1) energy production, resulting in an increase in acetogenesis, and 2) maintenance of redox balance, resulting in an increase in fermentative metabolism, lactate production, and conversion of NADH to NAD^+^. Increased levels of acetate compared to lactate under optimal aeration conditions suggests that demand for ATP is in excess of demand for NAD^+^.

Elevated respiratory activity of Δ*agr* is expected to increase endogenous ROS (4). To test this idea, we assessed ROS accumulation in bulk culture by flow cytometry of Δ*agr* and wild-type stains using carboxy-H2DCFDA, a dye that becomes fluorescent in the presence of several forms of ROS. As shown in Figure 6, ROS levels increased with *agr* deficiency, indicating correlation between *agr* activity, lower ROS levels, and increased bacterial survival in response to exogenous H_2_O_2_. These data help explain elevated lethality of peroxide in the absence of *agr*. Since lower ROS accumulation in wild-type cells correlates with decreased respiration and protection from killing by H_2_O_2_, the data also support the idea that suppression of endogenous ROS is key to *agr*-mediated protection from exogenous H_2_O_2_-mediated lethality.

**Figure 6.**
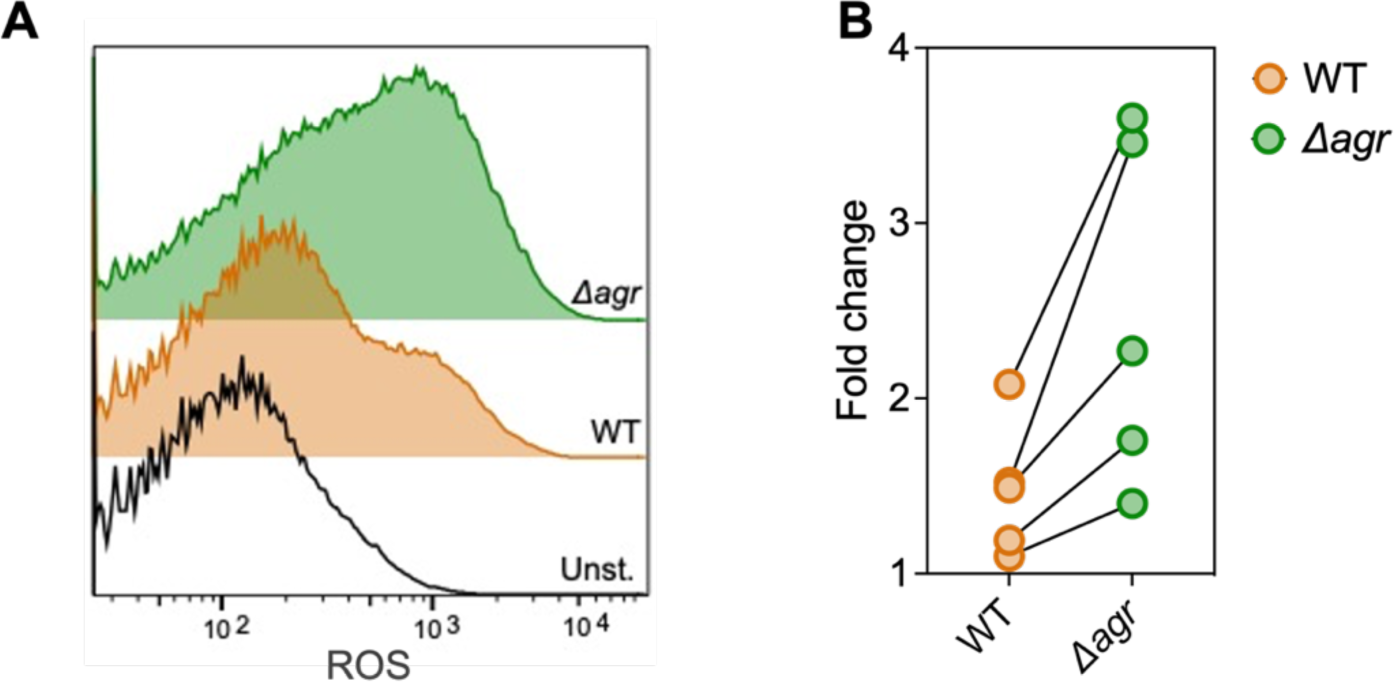
Increase in ROS levels associated with Δ*agr* deficiency. Flow cytometry measurements. *S. aureus* LAC wild-type (WT, BS819) and *Δagr* mutant (BS1348) were grown overnight, diluted, cultured in TSB medium for 1 h, and treated with carboxy-H2DCFDA (10 µM) for 5 min. Relative cell number is on the vertical axis. Unst. indicates samples containing LAC wild-type cells not treated with carboxy-H2DCFDA. (B) Five replicate experiments gave similar results (“fold change” indicates the mean wild-type or *Δagr* ROS level divided by the mean autofluorescence background signal; lines connect results in replicate experiments).

### Transcriptional changes due to Δagr mutation are long-lived and result in down-regulation of H_2_O_2_-stimulated genes relative to those in an agr wild-type

We reasoned that the transcriptional changes due to the Δ*agr* mutation likely persist, as does this strain’s susceptibility to killing by H_2_O_2_, after growth from overnight culture. With this in mind, and to determine whether *agr*-mediated changes act through *rot*, we performed RNA-seq experiments after 1-hr growth from overnight cultures of a Δ*agr* Δ*rot* double mutant that phenocopies wild-type with respect to H_2_O_2_-mediated death and with respect to its parental Δ*agr* strain (Supplementary file 3). Fold-changes and number of genes differentially expressed were lower in the Δ*agr* mutant relative to the wild-type culture, potentially because a significant portion of the population, even after an hour of growth (early exponential phase), still consisted of cells experiencing stationary phase at the time of sampling. Nevertheless, we did observe a shift to expression of fermentation-associated genes (*ilvA, pflAB*, *aldh1*, *ddh*, *lctp2*) in the Δ*agr* strain (Figure 4C and Supplementary file 3). Thus, up-regulation of metabolic genes in the Δ*agr* mutant extends beyond post-exponential growth to the exit from stationary phase and into subsequent cell proliferation, as does the long-lived protection from H_2_O_2_-mediated killing seen with the wild-type strain.

To examine induction of genes by lethal levels of H_2_O_2_, our gene expression analysis included a comparison between untreated and H_2_O_2_-treated cells after growth from overnight culture (Supplementary file 3). The Δ*agr* Δ*rot* double mutant that phenocopies wild-type had elevated expression of many genes involved in lowering oxidative stress compared to the Δ*agr* mutant. Those genes are involved in the regulation of misfolded proteins (*mcsA*, *mcsB*, *clpC*, *clpB*), Fe-S cluster repair (*iscS*), DNA protection and repair (*dps*), and genes regulated by the protein-damage repair gene *bshA (fhuB/G, queC-E)* (23) (Figure 7, Figure 7—figure supplement 1, and Supplementary file 3). Elevated expression of protective genes suggests that the double mutant survives damage from H_2_O_2_ better because protective genes are rendered inducible (loss of Rot-mediated repression). Overall, the data show that *agr* wild-type cells assume a long-lived stage after activation at high cell density in which they are primed to express genes (e.g., *clpB/C, dps*) that protect against high levels of exogenous oxidative stress.

**Figure 7.**
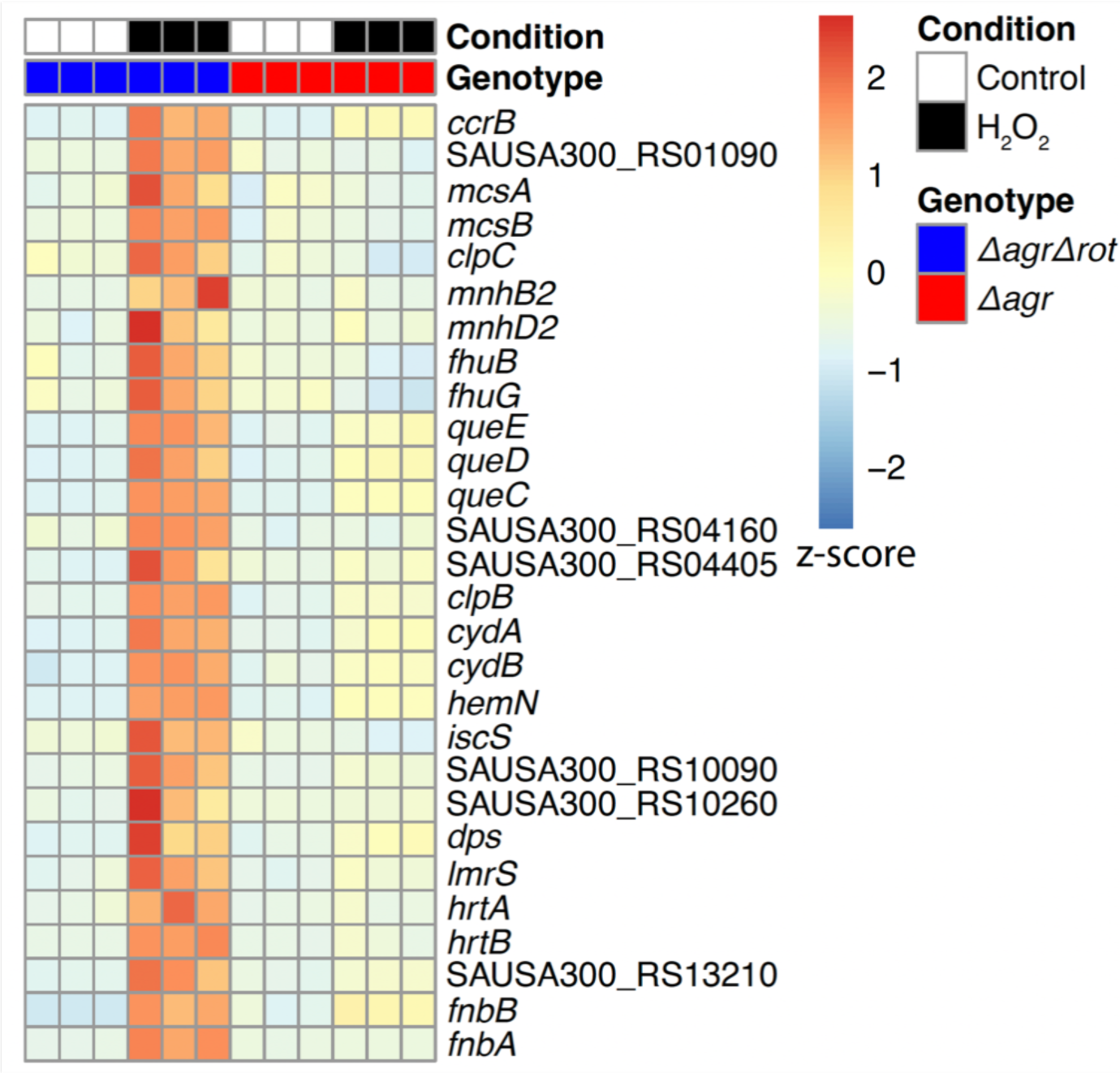
Rot-mediated up-regulation of H_2_O_2_-stimulated genes relative to those in an *agr* mutant. Genes shown are those up-regulated in a *Δagr Δrot* double mutant (BS1302) relative to that observed with the *Δagr* strain (BS1348). H_2_O_2_ treatment was for 30 min. Peroxide concentrations for *Δagr* (2.5 mM H_2_O_2_) and *Δagr Δrot* (10 mM H_2_O_2_) were determined to achieve ∼50% cell survival [see Methods and Figure 7—figure supplement 1]). RNA-seq data are from three independent cultures. Heatmap colors indicate expression z-scores. See Supplementary file 3 for supporting information.

### Endogenous ROS is involved in agr-mediated protection from lethal, exogenous H_2_O_2_ stress

We next monitored the effect of reducing respiration and ATP levels by adding subinhibitory doses of the redox cycling agent menadione (24) to cultures of Δ*agr* and wild-type cells prior to lethal levels of H_2_O_2_. Addition of menadione for 30 min, which induces a burst of ROS that inactivates the TCA cycle and thereby respiration (24), protected the Δ*agr* mutant but had little effect on the wild-type strain (Figure 8A). Menadione’s effect on respiration and ATP can be reversed by N-acetyl cysteine (24). Addition of N-acetyl cysteine in the presence of menadione restored H_2_O_2_ susceptibility to the *agr* mutant (Figure 8A). Thus, blocking endogenous ROS production/accumulation reverses the lethal effect of an *agr* deficiency with respect to a subsequent exogenous challenge with H_2_O_2_.

**Figure 8.**
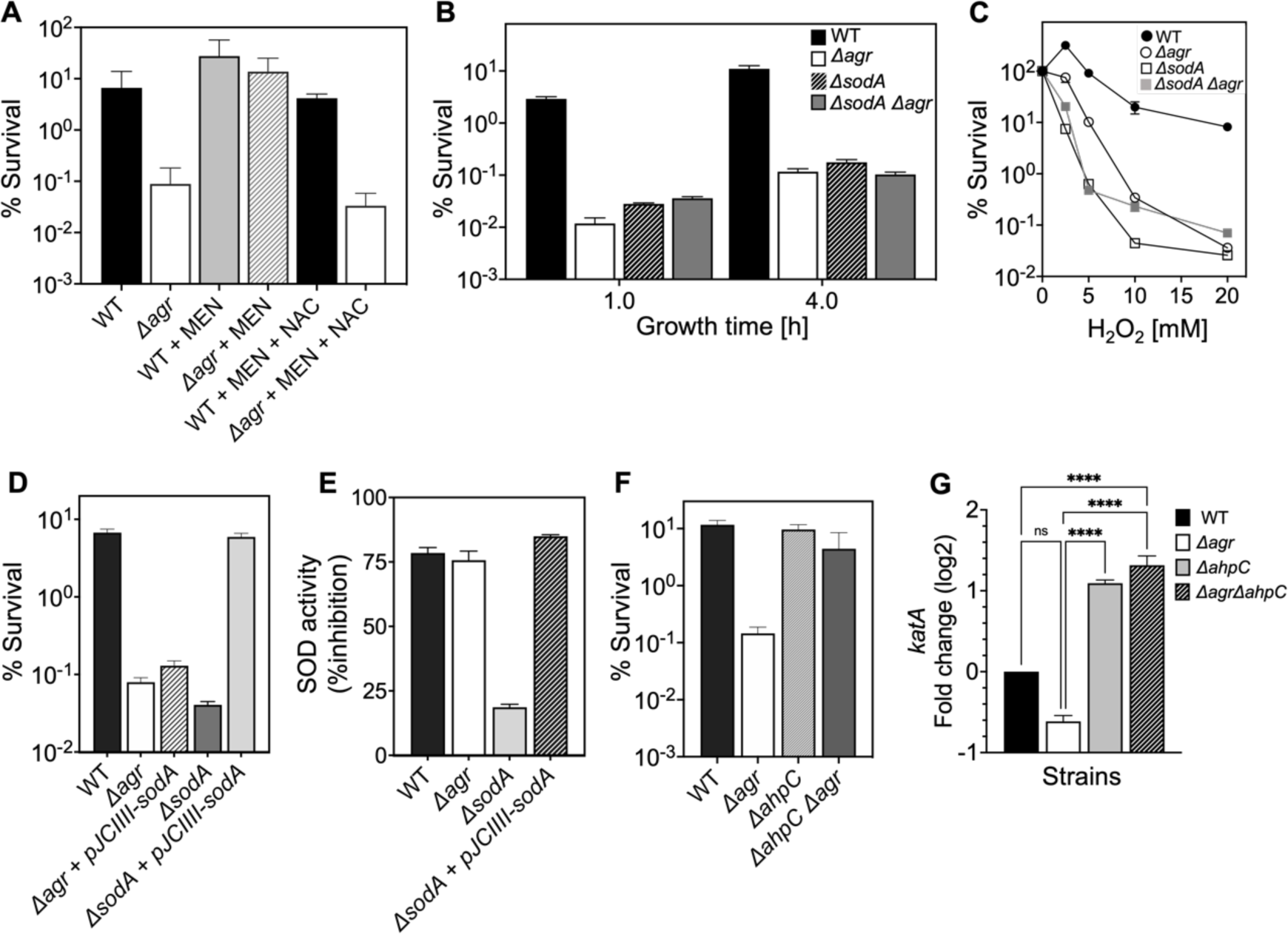
Involvement of endogenous ROS in *agr*-mediated protection from lethal H_2_O_2_ stress. (A) Protective effect of menadione on survival. *S. aureus* LAC wild type (BS819) and Δ*agr* mutant (BS1348) cultures were grown to late exponential phase (4 h after dilution of overnight cultures), exposed to 80 μM menadione (MD) with or without 4 mM N-acetyl cysteine (NAC) for 30 min prior to treatment with H_2_O_2_ (20 mM for 1 h) and measurement of survival. (B) Effect of *sodA* deletion on survival. Cultures of wild-type (BS819), Δ*agr* (BS1348), a *sodA::tetM* (BS1422)*, and sodA::tetM-agr* double mutant (BS1423) were grown to early (1 h after dilution, OD_600_∼0.15) or late log (4 h after dilution, OD_600_∼4.0) prior to treatment with 20 mM H_2_O_2_ for 60 min. (C) Effect of H_2_O_2_ concentration on survival. Late log (4 h, OD_600_∼4.0) cultures of the wild-type and Δ*agr* mutant strains were treated with indicated concentrations of H_2_O_2_ for 60 min. (D) Complementation of *sodA* deletion mutation. A plasmid-borne wild-type *sodA* gene was expressed under control of the *sarA* constitutive promoter (pJC1111-*sodA*) in late log-phase (4 h, OD_600_∼4.0) cells treated with 20 mM H_2_O_2_ for 60 min. (E) SodA activity. Wild-type or the indicated mutants were grown to late-exponential phase (OD_600_∼4.0); Sod activity was measured as in Methods. (F) Effect of *ahpC* deletion on survival. Late log-phase cultures of wild-type (BS819), *Δagr* (BS1348), *ahpC::bursa* (BS1486), and *ΔahpC::bursa*-*agr* double-mutant (BS1487) cells were treated with 20 mM H_2_O_2_ for 60 min. (G) Effect of *ahpC* deletion on expression of *katA* in the indicated mutants. Total cellular RNA was extracted from late exponential-phase cultures (OD_600_∼4.0), followed by reverse transcription and PCR amplification of the indicated genes, using *rpoB* as an internal standard. mRNA levels were normalized to those of each gene to wild-type control. Data represent the means ± S.D. from (*n* = 3) biological replicates. One-way ANOVA was used to determine statistical differences between samples (\*\*\*\**P* < 0.0001).

Rowe et al. (24) showed that menadione exerts its effects on endogenous ROS by inactivating the TCA cycle in *S. aureus*. To determine whether this mechanism can induce protection in the Δ*agr* mutant, we inactivated the TCA cycle gene *acnA* in *agr* wild-type and Δ*agr* strains (Figure 8—figure supplement 2). We found that *ΔacnA* mutation completely protected the Δ*agr* mutant from peroxide killing after growth to late exponential growth phase but had little effect on the wild-type *agr* strain. This finding supports the idea that TCA cycle activity contributes to an imbalance in endogenous ROS homeostasis in the Δ*agr* mutant, and that this shift is a critical factor for Δ*agr* hyperlethality. When we evaluated long-lived protection by comparing survival rates of *Δagr ΔacnA* mutant and *Δagr* cells following dilution of overnight cultures and regrowth prior to challenge with H_2_O_2_, *ΔacnA* remained protective, but less so (Figure 8—figure supplement 2). These partial effects of an *ΔacnA* deficiency suggest that *Δagr* stimulates long-lived lethality for peroxide through both TCA-dependent and TCA-independent pathways.

*S. aureus* has multiple enzymes that control the endogenous production and detoxification of ROS. SodA and SodM dismutate superoxide (O_2_^•-^) to H_2_O_2_, and catalase and AhpC then convert H_2_O_2_ to water, limiting the formation of toxic hydroxyl radical (OH^•^). Accordingly, we asked whether mutations in these pathways affect *agr*-dependent phenotypes with respect to lethal H_2_O_2_ exposure. A deficiency in the *sodA* superoxide dismutase (25) resulted in lower survival of the wild-type strain, similar to that observed with the Δ*agr* mutant (Figure 8B-C). The effect was reversed by complementation with *sodA* on a low-copy-number plasmid (Figure 8D). The Δ*sodA* mutation had no effect on killing with the Δ*agr* strain. Moreover, *sodA* expression (Supplementary file 1) and activity levels (Figure 8E) were similar for wild-type and the Δ*agr* mutant. Together, these observations suggest that the contribution of *sodA* toward protective priming by wild-type involves dismutation of low levels of endogenous superoxide generated by respiration. In contrast, endogenous levels of ROS are saturating for *sodA* in Δ*agr* cells. Inactivation of *sodM*, which is thought to be primarily induced by exogenous oxidative stress (26), had no noticeable effect on the H_2_O_2_ susceptibility of the wild-type or the Δ*agr* mutant. We conclude that scavenging enzymes, such as SodA, are better able to control the threat posed by endogenous ROS in wild-type than in Δ*agr* cells. They render the former better able to survive a subsequent lethal dose of H_2_O_2_, a compound that freely enters cells (27) and would add to endogenous ROS levels.

Other oxidative-stress-response mutations in genes encoding catalase, thiol-dependent peroxidases, and bacillithiol showed little effect on the relative lethality of H_2_O_2_ between wild-type and Δ*agr* mutant strains (Figure 8—figure supplement 1). Thus, protection against H_2_O_2_ lethality by these genes is not *agr*-specific. Paradoxically, a deficiency of *ahpC* (*ahpC::bursa*), which encodes a peroxidase (28), almost completely reversed the elevated killing associated with the Δ*agr* mutation (Figure 8F). An *ahpC* deficiency had no effect on the response of the otherwise wild-type strain. A deficiency in other downstream genes in the *ahpC* operon (*ahpF*, SAUSA300_0377-0378) showed no effect, indicating that the protective behavior of mutant *ahpC* was not caused by polar effects (Figure 8—figure supplement 3).

Results with *ahpC* deficiencies were initially surprising, because reduced ROS detoxification should increase rather than decrease killing. Compensatory expression of other protective genes, such as *katA* in the Δ*ahpC* Δ*agr* double mutant (28), might enable cells to better survive damage from subsequent stress-stimulated ROS increases. Indeed, *katA* expression increased > 10-fold in the Δ*ahpC* and Δ*ahpC* Δ*agr* double mutants (Figure 8G). Thus, Δ*katA* overcomes Δ*ahpC*-mediated protection, consistent with the idea expressed previously that *katA* is more protective than *ahpC* against high levels of exogenous oxidative stress (28, 29). We conclude that the protective action of an *ahpC*-deficient mutant is due to a pre-induced, compensatory increase in expression of another protective catalase.

### Importance of the long-lived “memory” of agr-mediated protection in a murine intraperitoneal infection model

To determine whether long-lived *agr*-mediated protection is important for *S. aureus* pathogenesis, we used the mixed infection strategy (outlined in Figure 3) in which a Δ*agrBD* mutant is “primed” in response to AIP produced by a Δ*rnaIII* mutant after overnight co-culture containing an equal ratio of the two mutant strains (Figure 9A and Figure 9— figure supplement 1). Then mice were infected via intraperitoneal inoculation; 2 h later, we lavaged the peritoneal cavity and harvested organs for determination of colony forming units (CFU). By 2 h after bacterial administration, the number of *S. aureus* cells injected as inoculum had declined by 1000-fold (Figure 9B and Figure 9—figure supplement 1). Mutant proportions, identified by differential plating, demonstrated that Δ*agrBD* cells were enriched by ∼30% in both peritoneum and organs compared to the Δ*rnaIII* mutant. The fraction of Δ*agrBD* (*rnaIII*^+^) mutants in sites of bacterial dissemination (heart, kidney, liver, lungs, and spleen) was similar to their elevated fraction in the peritoneum, thereby suggesting that *agr* enhances intraperitoneal infection and access to, rather than entry into extraperitoneal organs. In a control infection in which Δ*agrBD* was “unprimed” by mixing Δ*agrBD* and Δ*rnaIII* mutants immediately before growth from stationary phase, the proportion of Δ*agrBD* bacterial burden was lower at all tissue sites (Figure 9A and Figure 9—figure supplement 1). This drop represented a decline in long-lived *agr* induction of virulence.

**Figure 9.**
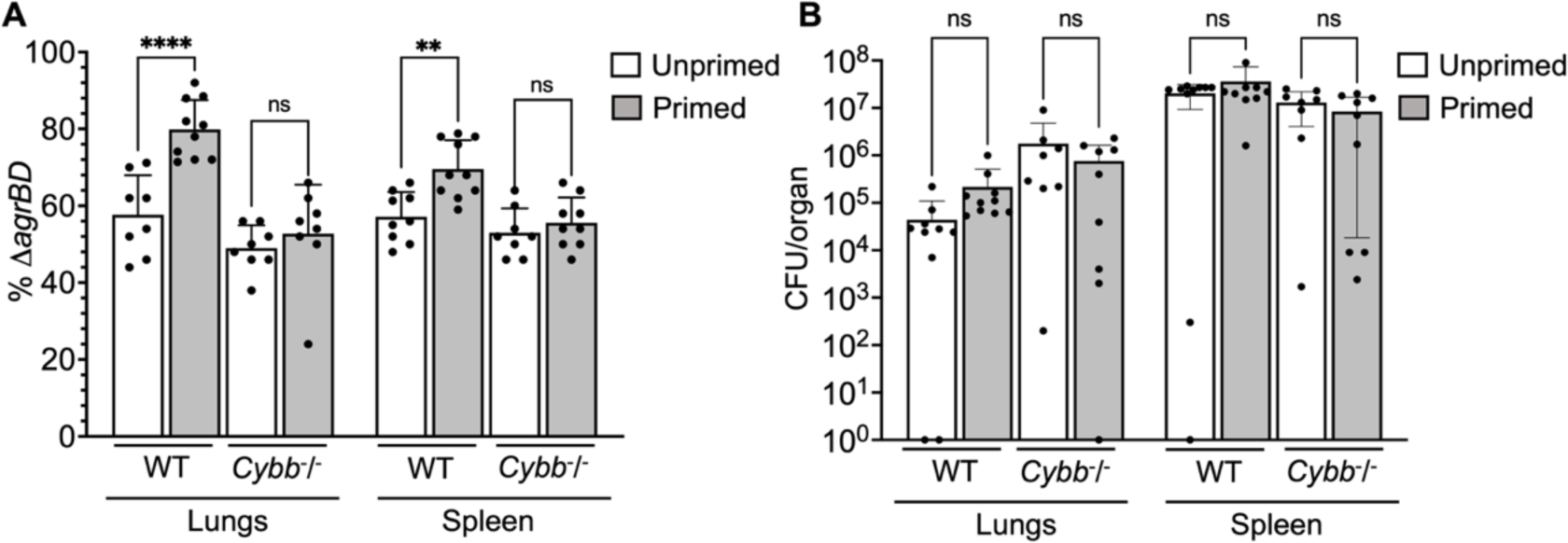
Survival advantage of *agr* priming of *S. aureus* absent in phagocyte NADPH-deficient murine infection. (A) percentage of 11*agrBD* (AIP-responsive in-frame deletion mutant carrying an intact RNAIII) cells and (B) bacterial burden in lung or spleen after 2 h of intraperitoneal infection of wild-type (WT) C57BL/6 mice or phagocyte NADPH oxidase-deficient (*Cybb^-/-^*) mice (see Figure 9—figure supplement 1 for data with other organs). *ΔagrBD* and *ΔrnaIII* mutant cultures were grown separately and mixed at a 1:1 ratio either before (primed) or after (unprimed) overnight growth, as for Figure 3. Both primed and unprimed mixtures were diluted after overnight growth, grown to early log phase (OD_600_∼0.15), and used as inocula (1 x 10^8^ CFU) for intraperitoneal infection (*n* = 2 groups of 10 mice each). After 2 h, lungs and spleen were harvested and homogenized; aliquots were diluted and plated to enumerate viable bacteria. Output ratios and total and mutant CFU from tissue homogenates were determined as for Fig 4E and 4H. A Mann-Whitney test (panel 9A) or Student’s two-tailed *t* test (panel 9B) were used to determine the statistical significance of the difference between primed and unprimed cultures. Error bars indicate standard deviation (\*\**P* < 0.01; \*\*\*\**P* < 0.0001).

To study *agr*-ROS effects during infection, we repeated *in vivo* studies using *Nox2*^−/−^ (also known as *Cybb*^−/−^) mice deficient in enzymes associated with host phagocyte production of ROS (the gp91 [phox] component of the phagocyte NADPH oxidase)(30). We found that *agr*-mediated priming (mixing Δ*agrBD* and Δ*rnaIII* before overnight co-culture) failed to increase hematogenous dissemination to lung and spleen tissues following infection of *Nox2*^−/−^ mice (Figure 9). Thus, when the host makes little ROS, long-lived *agr*-mediated protection has little effect in these tissues. The data also indicate that *agr*-mediated protection against ROS enhances fitness in lung and spleen, but it is dispensable for full virulence in other organs. Collectively, the murine experiments indicate that the long-lived “memory” of *agr* induction enhances overall pathogenicity of *S. aureus* during sepsis. They also support data previously published indicating that clearance of disseminating bloodstream pathogens (31) and protection from ROS buildup (32) are tuned to particular sites in the host organism.

## Discussion

We report that *agr*, a quorum-sensing regulator of virulence in *S. aureus*, provides surprisingly long-lived protection from the lethal action of exogenous H_2_O_2_. The protection, which is uncoupled from *agr* activation kinetics, arises in part from limiting the accumulation of endogenous ROS. This apparent tolerance to lethal stress derives from an RNAIII-*rot* regulatory connection that couples virulence-factor production to metabolism and thereby to levels of ROS. Collectively, the results suggest that *agr* anticipates and protects the bacterium from increases in ROS expected from the host during *S. aureus* infection.

Details of *agr*-mediated protection are sketched in Figure 10. At low levels of ROS, *agr* is activated by a redox sensor in AgrA, RNAIII is expressed and represses the Rot repressor, thereby rendering protective genes (e.g., *clpB/C, dps*) inducible via an unknown mechanism (induction, candidate protective gene(s), and their connection to endogenous ROS levels are being pursued, independent of the current report). Superoxide dismutase and scavenging catalases/peroxidases detoxify superoxide and peroxide, respectively (scavenging deficiencies reduce the protective effect of wild-type). Deletion of *agr* eliminates expression of RNAIII and repression of Rot, resulting in a metabolic instability associated with a 100-fold increase in H_2_O_2_-mediated death.

**Figure 10.**
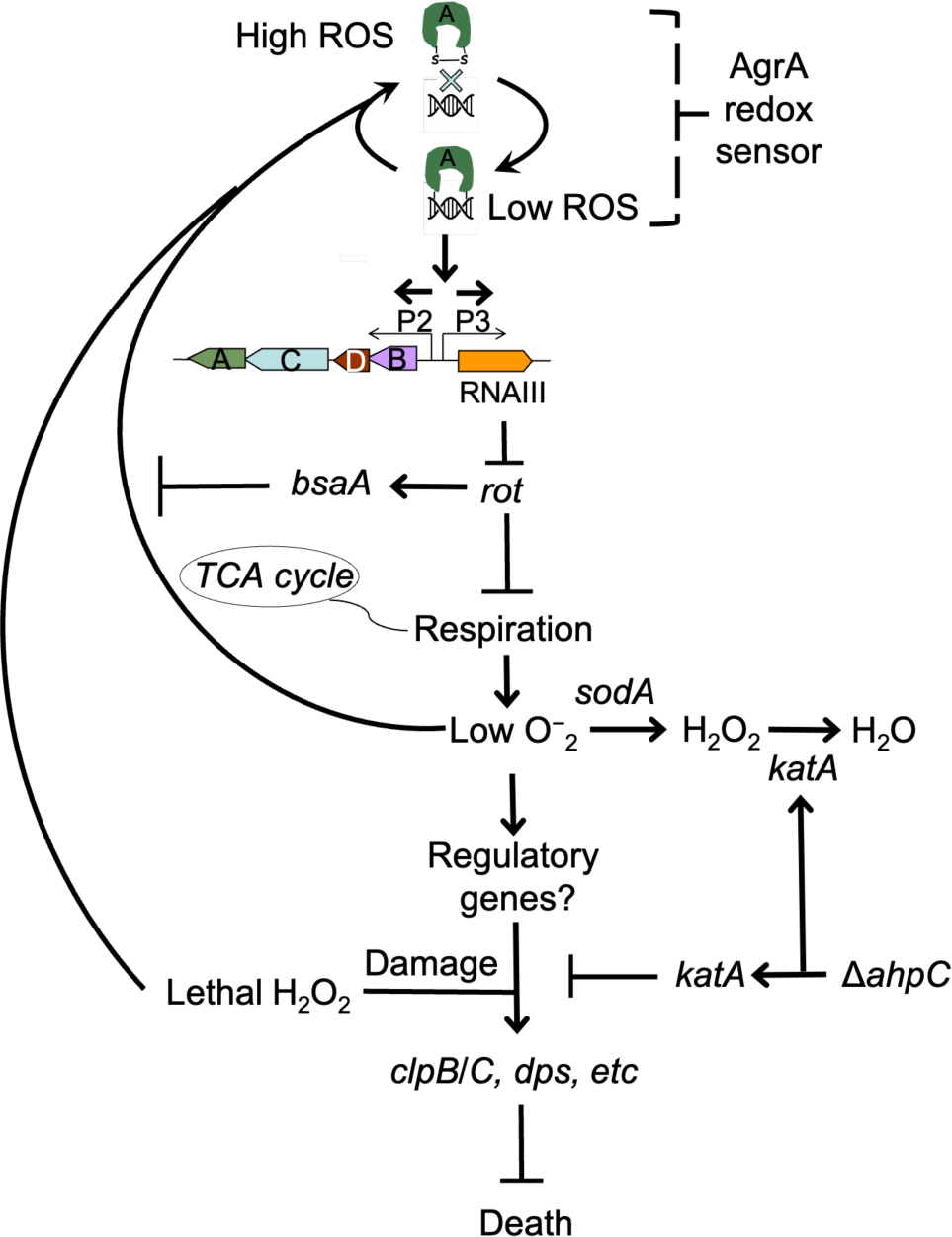
Schematic representation of *agr*-mediated protection from ROS. At low levels of oxidative stress, the redox sensor in AgrA binds to DNA at promoters P2 and P3, activating expression of the two operons. Expression of RNAIII blocks translation of Rot, which decreases respiration and production of superoxide. ROS quenchers (*sodA* and *katA*/*ahpC*) suppress formation of most ROS that would otherwise signal the redox sensor in AgrA to halt stimulation of RNAIII expression and the production of further superoxide via respiration. This feedback system regulates respiration thereby limiting the accumulation of ROS in wild-type cells. Wild-type cells are primed for induction of protective genes (e.g., *clpB/C, dps*) by loss of the *rot* repressor system via an unknown mechanism when cells experience damage from high levels of oxidative stress (experimentally introduced as lethal exogenous H_2_O_2_); *Δagr* cells that experience high levels of endogenous H_2_O_2_ fail to induce protective genes. Exogenous H_2_O_2_ or high levels of endogenous ROS, for example from extreme stress due to ciprofloxacin (12), lower RNAIII expression and allow Rot to stimulate *bsaA* expression, which produces a protective antioxidant. The protective action of an *ahpC* deficiency acts through compensatory expression of *katA*, which results in more effective scavenging of H_2_O_2_ produced from increased respiration in Δ*agr* strains and/or exogenous lethal H_2_O_2_.

The *agr* system directly reduces H_2_O_2_-mediated killing by reducing levels of endogenous ROS, much like intrinsic tolerance to lethal antimicrobial stress (33). However, the protective system we describe is distinct in that it primes cells for induction of genes (e.g., *clpB/C, dps*) that mitigate damage upon subsequent exposure to high levels of ROS. Still unidentified protective genes exist; thus, *agr-*mediated protection may be further shaped by both known (*ahpC*) and unidentified pathways and factors that modify the redox state. Another distinctive feature of *agr*-mediated protection is its manifestation even in early log-phase cultures, long after the maximal transcription of *agr* at high cell density, i.e., quorum. In a sense, *S. aureus* has a “memory” of the *agr*-activated state.

Transcriptional profiling during growth from diluted, overnight cultures revealed that the Δ*agr* mutation elevated expression of several respiration and fermentation genes. Acceleration of cellular respiration is likely a source of ROS, as it appears to be for bactericidal antibiotics (3). Our work supports this idea by showing that increased respiration caused by deletion of *agr* is associated with increased ROS-mediated lethality. How *agr* deficiency is connected to the corruption of downstream processes that result in metabolic inefficiency and increased endogenous ROS levels is unknown. Given that Δ*agr* mutants are unable to downregulate surface proteins during stationary phase (34, 35), it is possible that deletion of *agr* perturbs the cytoplasmic membrane or the machinery that sorts proteins across the cell wall. In support of this notion, jamming SecY translocation machinery of *E. coli* results in downstream events shared with antibiotic lethality, including accelerated respiration and accumulation of ROS (36). In this scenario, the formation of a futile macromolecular cycle may accelerate cellular respiration to meet the metabolic demand of unresolvable problems caused by elevated surface sorting.

As noted above, *agr* is inactivated by oxidation, which elevates levels of the antioxidant BsaA during exposure to H_2_O_2_ (7). That would make our finding that H_2_O_2_-mediated killing is increased in the *Δagr* mutant paradoxical. This apparent inconsistency can be explained by a focus of prior work on growth-related phenotypes (7) rather than on lethality (the underlying mechanisms are distinct (2)). Additionally, we note that *bsaA* was not upregulated in either our RNA-seq experiments (Supplementary file 1) or in previous transcriptional profiling data (37). Thus, an alternative, but not mutually exclusive, hypothesis is that the growth-related effect of *bsaA* on *agr*-mediated responses to stress is strain dependent. Another complexity involves test conditions, as indicated by consideration of previous work in which wild-type cells exhibited greater oxidative stress than the *agr*-deficient mutant due to *agrA*-mediated production of ROS-inducing phenol-soluble modulins (37). The present experiments were performed in highly diluted cultures in which levels of these modulins are likely low (38, 39). The complex relationship between *agr*, ROS-mediated lethality, and physiological state illustrates the importance of understanding *agr* biology before applying therapies that inactivate *agr* (8).

We also note that although the absence of *agr* increases killing by high levels of H_2_O_2_, it has the opposite effect for lethal concentrations of ciprofloxacin (12). In the latter case, the absence of *agr* upregulates expression of *bsaA* in the strain examined; *bsaA* counters endogenous ROS induced by ciprofloxacin (12). The present work shows that excess endogenous ROS is generated during *agr* deficiency. Thus, protection from endogenous lethal stress via *agr* inactivation may not be only through the redox-dependent *bsaA* but also by a second pathway involving increased respiratory metabolism. The present work also supports the idea that exposing bacteria to exogenous H_2_O_2_ does not fully recapitulate the intracellular environment created by antibiotics and other stresses that act via ROS-mediated cell death (36), emphasizing that inactivation of *agr* can be either destructive or protective, depending on the type of lethal stress. Similar results have been reported with other bacteria: *mazF*, *lepA*, and *cpx* are destructive or protective based on the level of lethal stress (40, 41).

The protective activity of *agr* carried over to *in vivo* studies using mice, as it was largely absent if the mice were deficient in host phagocyte production of ROS (*Nox2*^−/−^ mice with a null allele of NADPH oxidase). The benefits of *agr* to *S. aureus* fitness seen with NADPH oxidase-proficient mice were observed largely in lungs, a key host defense niche for neutrophil-mediated clearance of disseminating pathogens (31). The redox switch in AgrA, plus the protective properties associated with *agr* activation, lead to a clinical model in which *agr* links virulence-factor expression to an intrinsic protection against a lethal, H_2_O_2_-mediated immune response during infection (Figure 11). In this model, *agr* quorum-sensing renders cells better prepared to respond to lethal, exogenous oxidative stress. We note, however, that *agr*-mediated fitness benefits were present in certain tissues even in NADPH oxidase-deficient mice, indicating the existence of long-lived factors other than those that suppress oxidative stress. Thus, such a pre-emptive defense system may apply to many challenges experienced by *S. aureus* during infection, especially during bloodstream dissemination and conditions within inflamed tissues (42, 43).

**Figure 11.**
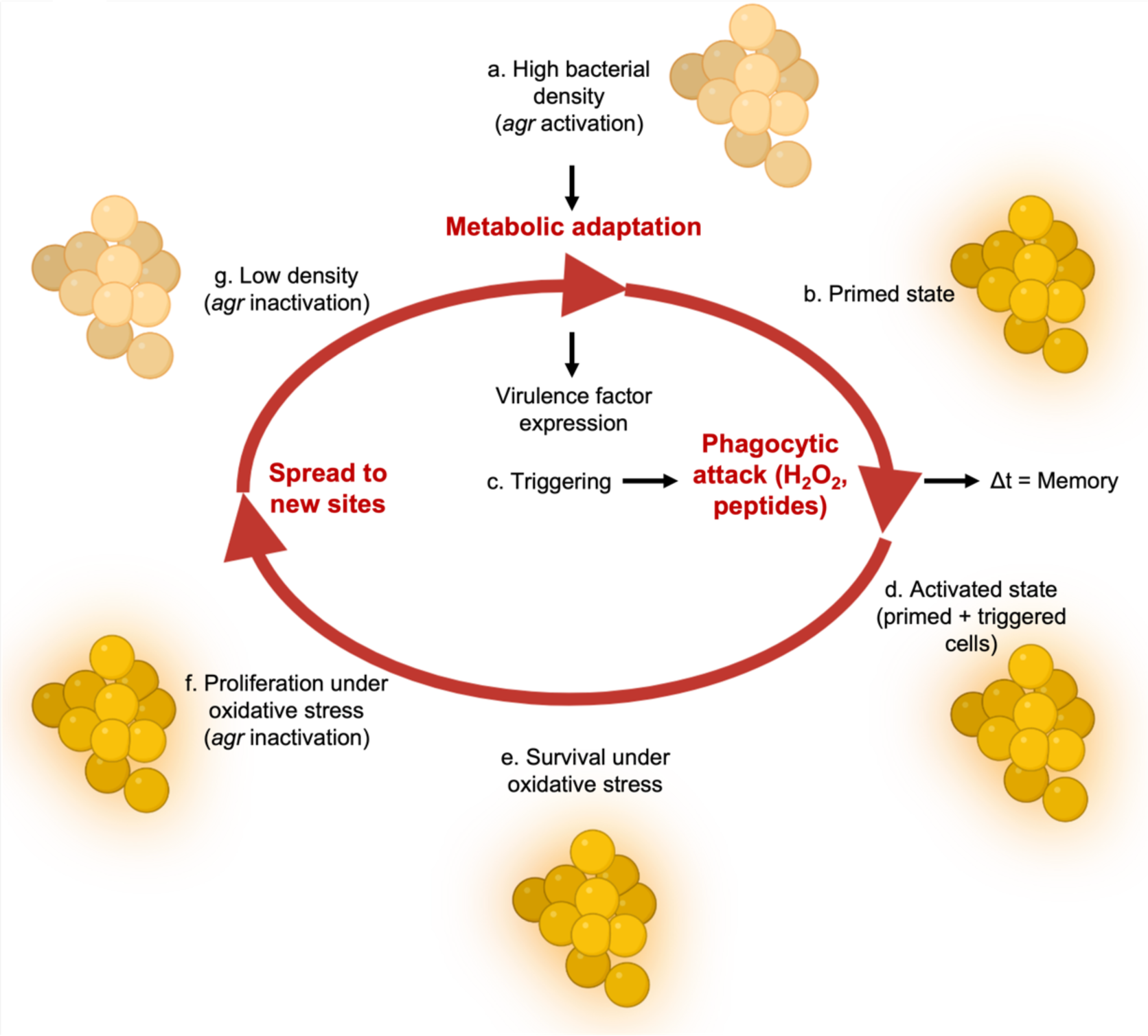
Relationship of *agr* priming and virulence. The ecology of abscess formation and subsequent bacterial dissemination can be described as a cycle. (a) During abscess formation, a hallmark of *S. aureus* disease, *agr* is activated by high bacterial cell density (quorum sensing) (63). (b) The bacterium assumes a primed stage due to repression of the *rot* repressor. (c, d, e) The lethal effects of immune challenge, which is called triggering (64), are survived by the persistence (“memory”) of the *agr*-activated state. (f) *agr* expression is inactivated by oxidation, thereby elevating expression of the antioxidant *bsaA* (7), which enables proliferation when oxidative stress is sublethal (7). (g) By surviving damage caused by lethal exogenous oxidative stress, primed *S. aureus* escape from the localized abscess to produce new infectious lesions (bloodstream dissemination) or to infect new hosts, where the cycle would be repeated.

In conclusion, uncoupling of *agr*-mediated tolerance from bacterial population density anticipates increases in exogenous ROS expected during *S. aureus*-host interactions, thereby contributing to virulence. The ubiquity of quorum sensing suggests that it protects many bacterial species from oxidative damage. A next step is to find RNA, protein, and/or epigenetic markers underlying the *agr*-mediated “memory” that improves protection against subsequent H_2_O_2_ exposure, since that will provide insights into the role of *agr* in cellular survival and adaptation during infection. Discovering ways to manipulate the lethal stress response, as seen with supplementation of antimicrobials with N-acetyl-cysteine during treatment of *Mycobacterium tuberculosis* (44) and development of inhibitors of enzymes that produce protective H_2_S (5), could reveal novel approaches for enhancing antimicrobial therapy and host defense systems (45–47).

## Materials and Methods

### Bacterial strains, plasmids, primers and growth conditions

*S. aureus* strains, plasmids, and primers used in the study are described in Tables 1 and 2. Bacterial cells were grown in Tryptic Soy Broth (TSB, glucose concentration at 2.5 g/L) at 37°C with rotary shaking at 180 rpm. For suboptimal aeration, broth cultures were grown in a closed-capped 15 mL conical tube with 10 mL of TSB. Colony formation was on Tryptic Soy Agar (TSA) with or without defibrinated sheep blood, incubated at 37°C or 30°C. Phages 80α and Φ11-mediated transduction was used for strain construction (48); transductants were selected on TSA plates containing the appropriate antimicrobial.

**Table 1.**
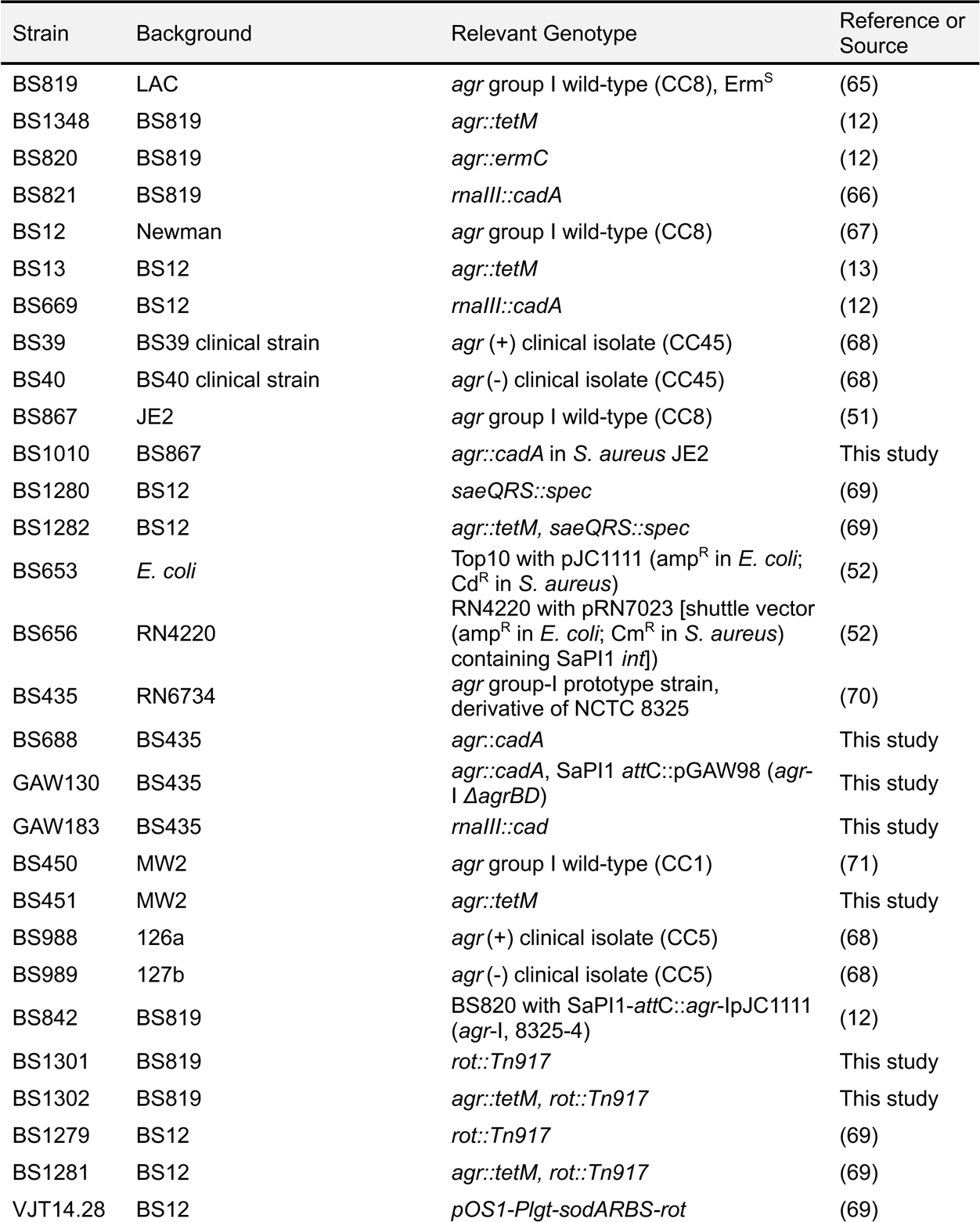
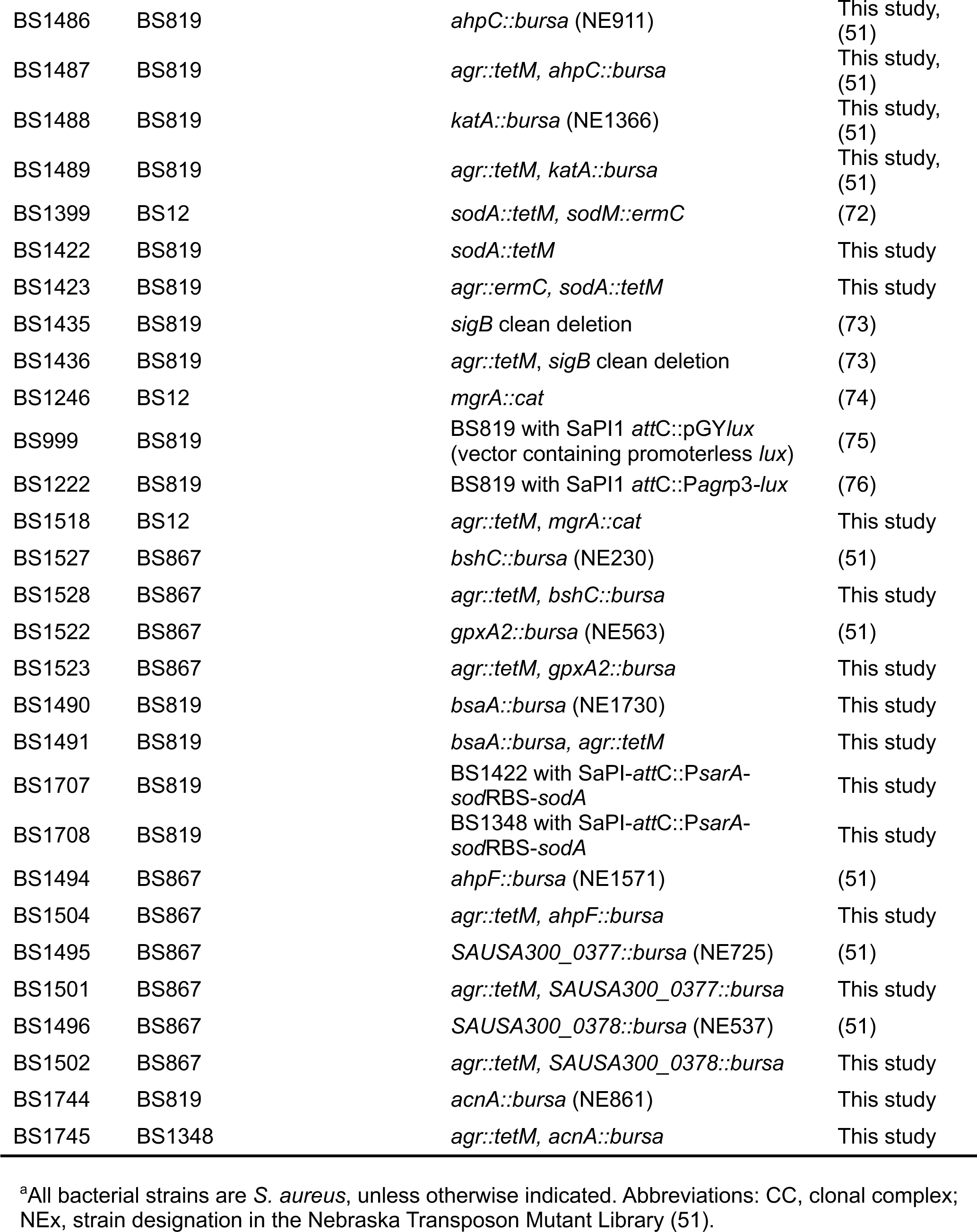
Bacterial strains^a^.

**Table 2.**
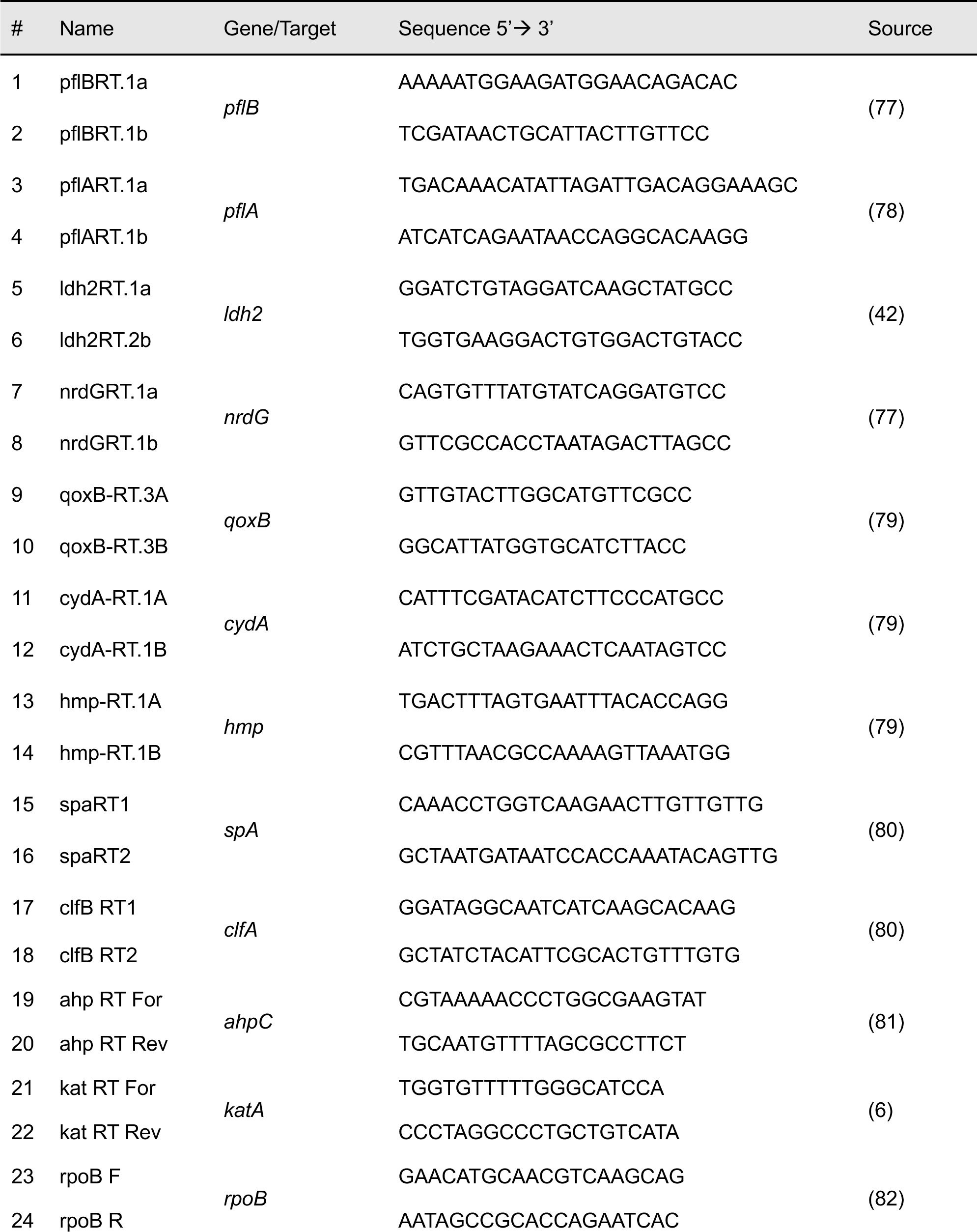
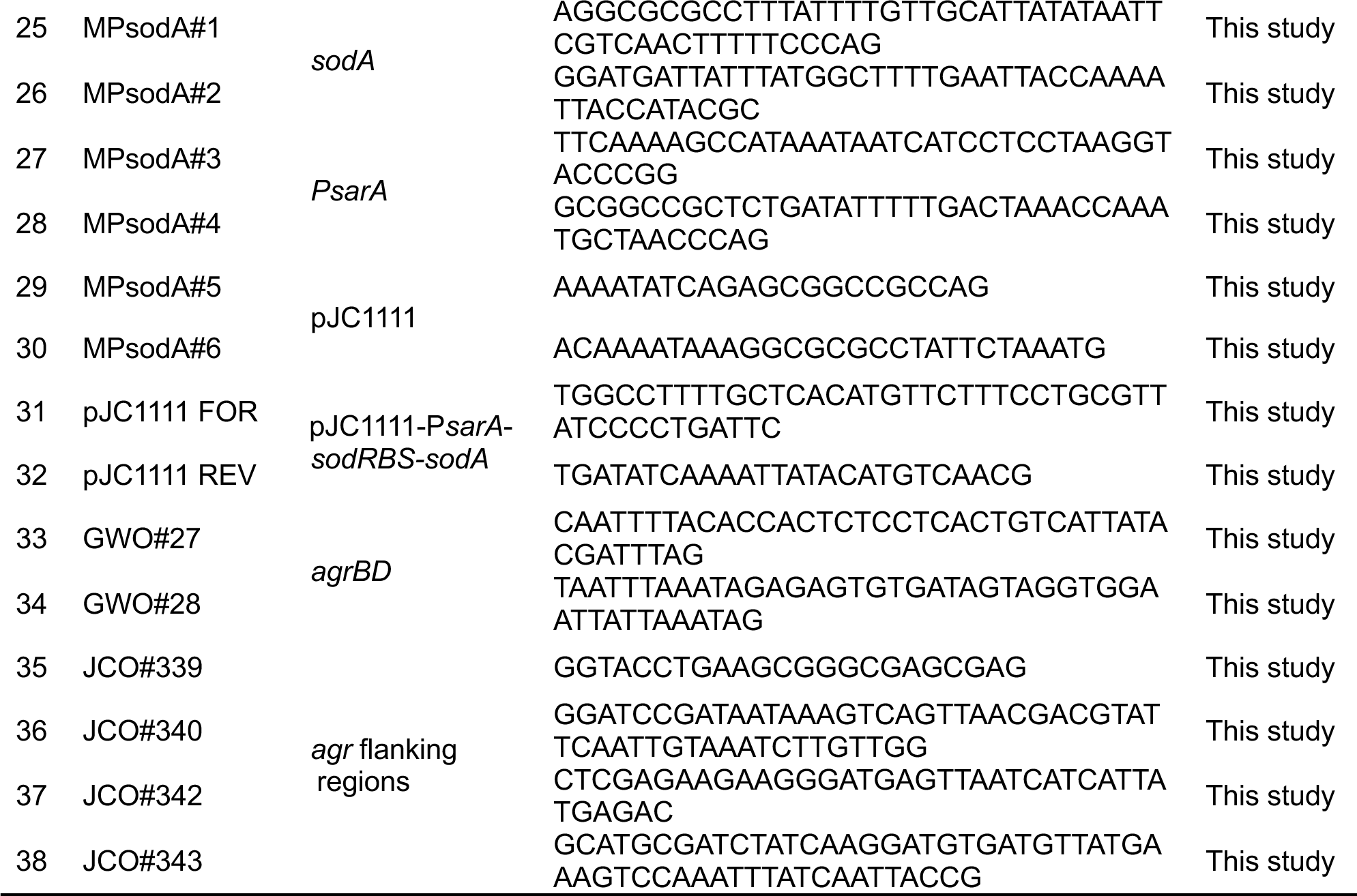
Oligonucleotides.

For analysis of *in vitro* growth curves, overnight cultures grown in TSB were diluted 1:1000 in chemically defined medium (CDM) (49), and growth was monitored at 37°C in 96-well plates (100 µL/well) using an Agilent LogPhase 600 Microbiology Reader (Santa Clara, CA) with 1 mm orbital shaking, measuring OD_600_ at 40-min intervals. The curves represent averaged values from five biological replicates. The exponential phase was used to determine growth rate (μ) from two datapoints, OD_1_ and OD_2_ flanking the linear portion of the growth curve, following the equation lnOD_2_-lnOD_1_/t_2_-t_1_, as described (50).

### Measurement of bioluminescence

Overnight cultures were diluted to OD_600_ ∼0.05 and grown in TSB at 37°C with rotary shaking at 180 rpm. Aliquots (100 μL) were inoculated into flat bottom 96-well microtiter plates (Corning, Corning, NY), and bioluminescence was detected using a BioTek Synergy Neo2 plate reader (Agilent, Santa Clara, CA).

### Antimicrobials and chemicals

Antimicrobials, chemicals, and reagents were obtained from MilliporeSigma (Burlington, MA) or Thermo Fisher Scientific (Waltham, MA).

### Construction of mutants

Transposon mutants were generated by transducing *Bursa aurealis* insertions, obtained from the University of Nebraska transposon mutant (ΦNE) library (51), into LAC or LAC *agr::tetM* using phages 80α and Φ11.

Construction of the Δ*agrBD* mutant: *S. aureus* Δ*agrBD* mutant GAW128 was generated by a chromosomal integration strategy outlined in Chen *et al*. (52) in an *agr*-null background strain, BS687. Plasmid pJC1111, a suicide plasmid containing a cadmium resistance (Cd^R^) cassette and the SaPI1 *attS* site that enables single-copy insertion into the corresponding chromosomal SaPI1 *att*C site, was used as the backbone vector for the *S. aureus agrBD* construct. pJC1000 contains the RN6734 *agr* locus cloned into pUC18. Inverse PCR of pJC1000 was performed using *agrBD* primers GWO#27 and GWO#28, re-ligated following treatment with polynucleotide kinase, and designated pGAW98. The *Sph*I-*EcoR*I fragment of pGAW98 was ligated into the SaPI1 integration vector pJC1111 and designated pGAW119. Strain RN9011 (RN4220 with pRN7023 [vector (Cm^R^) containing SaPI1 integrase]) was electroporated with plasmid pGAW119 and plated on GL agar containing 0.1 mM CdCl_2_. Phage 80α lysates of Cd^R^ colonies were used to transduce BS687 (RN6734 Δ*agr*::*ermC*, Erm), generating GAW128 (Δ*agrBD*).

To construct *agr* mutant BS687*, agr* flanking regions were amplified with primer pairs JCO#339, JCO#340, and JCO#342, JCO#343 and cloned into the *Hinc*II site of pUC18 to generate pJC1527 and pJC1528, respectively. pJC1530 was generated by four-way ligation of the *Kpn*I*-BamH*I fragment of pJC1527 (*agr* left flank), *Xho*I*-Sph*I fragment of pJC1528 (*agr* right flank), and *BamH*I-*Xho*I fragment from pJC1073 (Erm cassette) to *Kpn*I-*Sph*I digested pJC1202 (replacement vector). Plasmid pJC1530 was electroporated into strain RN4220 with selection on GL agar containing 10 µg/mL of chloramphenicol at 30°C. Phage 80α lysates of Cm^R^ colonies were used to transduce strain JCSA18 (*rpsL*\* mutant of RN6734 that results in streptomycin resistance) and then allelic exchange of the Em^R^ Sm^S^ Cm^R^ colonies was performed as previously described. Phage 80a lysates of Em^R^ Sm^R^ Cm^S^ colonies were then used to transduce RN6734 with selection for Em^R^, generating BS687.

*sodA* complementation: Plasmid PsarA-*sodA*-pJC1111, expressing *sodA* under the control of the constitutive promoter P*sarA*, was integrated into the *S. aureus* chromosome at the SaPI1 *attC* site of strain LAC (53), LAC *sodA::tetM*, and LAC *agr::ermC*. Complementation plasmid P*sarA*-*sod*RBS-*sodA* was generated by Gibson assembly and inserted into the SaPI1 integration vector pJC1111. Wild-type *sodA* and the *sarA* promoter were amplified from *S. aureus* gDNA using primers MPsodA#1-2 (*sodA* gene and RBS) and MPsodA#3-4 (P*sarA*). Primers MPsodA#5-6 were used to linearize pJC1111. Primers introduced relevant oligonucleotide overlaps that enabled Gibson assembly (6), generating P*sarA*-*sod*RBS-*sodA*. P*sarA*-*sod*RBS-*sodA* was transformed into *E. coli* DH5α for amplification, purification, and sequence validation via primers pJC1111 FOR and pJC1111 REV. Purified P*sarA*-*sod*RBS-*sodA* was electroporated into RN9011 and positive chromosomal integrants at the SaPI1 chromosomal attachment (*att*C) site were selected with 0.1 mM CdCl_2_. Phage 80a lysates of positive integrants were used to transduce BS1422 (LAC *sod*::*tetM*) and BS1348 (LAC *agr*::*tetM*), generating BS1707 and BS1708, respectively.

### Survival measurements

To measure lethal action, overnight cultures were diluted (OD_600_∼0.05) in fresh medium and grown with shaking to early exponential (OD_600_∼0.15) or late log (OD_600_∼4) phase, conditions when *agr* expression is largely absent (12) or maximally activated, respectively. Early (undiluted) and late exponential phase cultures (diluted into fresh TSB medium to OD_600_∼0.15) were incubated with H_2_O_2_ under aerobic conditions either at a fixed concentration for one or more time points or at various concentrations for a fixed time. At the end of treatment, aliquots were removed, concentrated by centrifugation and serially diluted in phosphate-buffered saline to remove H_2_O_2_, and plated for determination of viable counts at 24 h. Percentage of survival was calculated relative to a sample taken at the time of H_2_O_2_ addition. When menadione and N-acetylcysteine were used to inhibit or potentiate killing by H_2_O_2_, they were added prior to lethal treatments as described previously (14). For experiments involving menadione pretreatment, cultures were grown for 3.5 h, and menadione (40 mM solution in 96% EtOH, final concentration 80 μM) was added for the last 0.5 h of culture, preceding the H_2_O_2_ treatment at 4 h. N-acetylcysteine was used to counter the action of menadione; it was added simultaneously with menadione, at a final concentration of 30 mM (640 mM stock in sterile ddH_2_O was used). All experiments were repeated at least three times; similar results were obtained from the biological replicates.

### Measurement of glucose consumption

Overnight cultures were diluted into fresh TSB (OD_600_∼0.05) and grown for 4 hours with shaking at 180 rpm (OD_600_∼4) at 37°C. Glucose was assayed in the supernatant fluids of bacterial cultures following centrifugation at 12,000 x g, using Centricon-10 concentrators (MilliporeSigma, Burlington, MA), and pH adjustment to 6.5-7.0 using NaOH. Cells were assayed and plated hourly for determination of viable counts as indicated in figures. Glucose content was measured from serial dilutions of supernatants using UV method (cat. no. 10-716-251-035) following manufacturer’s instructions (R-Biopharm, Darmstadt, Germany). Glucose consumption was expressed as μg of glucose consumed over 3 h of culture per 10^8^ bacterial cells. There was no detectable glucose in culture supernatants at 4 h of culture (data not shown).

### Measurement of excreted metabolites

Excreted metabolites were assayed in the supernatant fluids of bacterial cultures following centrifugation at 12,000 x g for 10 min for late exponential (4 h, OD_600_∼4) or multiple time points (acetate), as indicated in figures. Aliquoted supernatants were stored at −80°C and thawed on ice prior to analysis. Cells were plated for determination of viable counts; L(+)-lactate and acetate concentrations were measured using commercially available colorimetric and fluorometric kits (cat. no. MAK065, ab204719), according to manufacturer’s recommendations (MilliporeSigma, Burlington, MA and Abcam, Cambridge, UK, respectively).

### Measurement of intracellular metabolites

Overnight cultures were diluted into fresh TSB (OD_600_∼0.05), grown for 4 hours at 37°C with shaking at 180 rpm (OD_600_∼4), and plated for determination of viable counts at 4 h. The remaining cells were concentrated by centrifugation at 12,000 x g for 10 min, and resuspended in lysis buffer provided by the assay kit. Cells were lysed by repeated homogenization (2 cycles of 45 sec homogenization time at 6 M/s followed by a 5 min pause on ice) using Lysing Matrix B tubes in a FastPrep-24 homogenizer (MP Biomedicals, Irvine, CA). After lysis, cell debris was removed by centrifugation (12,000 × *g*, 10 min) and the supernatant was used for determination of pyruvate, fumarate, citrate and acetyl-CoA levels using colorimetric (pyruvate, fumarate) or flurometric (citrate, acetyl-CoA) assays (cat. no. KA1674, ab102516, KA3791 and MAK039, respectively) and a microplate reader (BioTek Synergy Neo2, Agilent, Santa Clara, CA) according to manufacturer’s instructions (Abnova, Taipei City, Taiwan and Abcam, Cambridge, UK and MilliporeSigma, Burlington, MA, respectively). Assayed metabolites were measured in μg and normalized to cell count.

### Measurement of oxygen consumption

Overnight cultures were diluted into fresh TSB (OD_600_∼0.05), grown for 5 hours at 37°C with shaking at 180 rpm (OD_600_∼4), diluted (OD_600_∼0.025) in fresh TSB, and added to a microtiter plate (200 μL/well). Oxygen consumption rate (OCR) was measured using a Seahorse XF HS Mini Analyzer (Agilent, Santa Clara, CA) according to the manufacturer’s instructions. The Seahorse XF sensor cartridge was hydrated in a non-CO_2_ 37°C incubator with sterile water (overnight) and pre-warmed XF calibrant for 1 hour prior to measurement. OCR measurements were recorded in 15 measurement cycles with 3 minutes of measurement and 3 minutes of mixing per cycle. CFU were enumerated to confirm equal concentrations of *agr*-deficient mutant and wild-type cells

### Measurement of ATP, NAD^+^, and NADH

For ATP, overnight cultures were diluted into fresh TSB (OD_600_∼0.05), grown for 4 hours at 37°C with shaking at 180 rpm (OD_600_∼4), diluted (OD_600_∼1.0) in fresh TSB, and incubated at room temperature with reagent for determination of ATP using BacTiter-Glo Microbial Cell Viability Assay (cat. no. G8232; Promega, Madison, WI), according to the manufacturer’s instructions. Luminescence was detected a BioTek Synergy Neo2 plate reader (Agilent, Santa Clara, CA). The amount of ATP was calculated and normalized to cell count.

For NAD^+^ and NADH, overnight cultures were diluted into fresh TSB (OD_600_∼0.05), grown for 4 hours at 37°C with shaking at 180 rpm (OD_600_∼4), and plated for viable counts at 4h or concentrated by centrifugation at 12,000 x g for 10 min and resuspended in lysis buffer provided by the assay kit. Cells were lysed by repeated homogenization (2 cycles of 45 sec homogenization time at 6 M/s followed by a 5 min pause on ice) using Lysing Matrix B tubes in a FastPrep-24 homogenizer (MP Biomedicals, Irvine, CA). After lysis, cell debris was removed by centrifugation (12,000 × *g*, 10 min) and the supernatant was used for determination of NAD^+^ and NADH levels using a colorimetric assay kit (cat. no. KA1657; Abnova, Taipei City, Taiwan) and microplate reader (BioTek Synergy Neo2, Agilent, Santa Clara, CA) according to the manufacturer’s instructions.

### Measurement of baseline ROS levels

Overnight cultures were diluted into fresh TSB (OD_600_∼0.05), and grown with shaking to early exponential phase (OD_600_∼0.2). 200 µL of culture was removed and cell density was normalized before staining with carboxy-H2DCFDA fluorescent dye (final concentration 10 µM) (Invitrogen, Waltham, MA). Samples were incubated at room temperature for 5 minutes, then 800 µL of PBS + EDTA buffer (100 mM) was added to each sample, and ROS levels were measured by fluorescence-based flow cytometry (BD Fortessa, BD Biosciences, San Jose, CA). All tubes with cultures were wrapped with aluminum foil to avoid light. A sample containing LAC wild-type cells lacking carboxy-H2DCFDA was included as a control for auto-fluorescence. Forward and side scatter parameters were acquired with logarithmic amplification. ROS was detected using the 488 laser and a 530/30nm bandpass filter. Data were analyzed using FlowJo software version 10.8.1 (BD Biosciences, San Jose, CA).

### Measurement of superoxide dismutase (SOD) activity

Overnight cultures were diluted (OD_600_∼0.05) into fresh TSB, grown to late exponential phase (OD_600_∼4), diluted to OD_600_=1, centrifuged at 12,000 x g for 5 min, and the cell pellet was homogenized in 300 μL of ice-cold lysis buffer (100 mM Tris-HCl pH 7.4 + 0.5% Triton + 5 mM 2-mercaptoethanol + 0.2 mM PMSF). SOD activity was measured using a commercially available kit (cat. no CS0009-1KT), according to manufacturers’ instructions (MilliporeSigma, Burlington, MA). The experiment was repeated three times with similar results.

### RNA sequencing and data analysis

Overnight cultures were diluted (OD_600_∼0.05) into fresh TSB medium and grown at 37 °C to early exponential phase (OD_600_∼0.5) (*Δagr* single mutant and *Δagr Δrot* double mutant) or late exponential phase (OD_600_∼4) (wild-type and *Δagr* strains). Samples of *Δagr* and *Δagr Δrot* were divided into two 3 mL aliquots, and the aliquots were incubated at 37 °C for another 30 min, with or without treatment with H_2_O_2_. Peroxide concentrations for *Δagr* and *Δagr Δrot* were normalized to expected killing at the time of harvest (Figure 7—figure supplement 1).

Three independent cultures for each sample were used for determination of transcriptional profiles. Briefly, cultures were concentrated by centrifugation (12,000 x g for 5 min), and resuspended cells were disrupted using Lysing Matrix B tubes in a FastPrep-24 homogenizer (MP Biomedicals, Irvine, CA) at 6 M/s, for 30s, 3 times (samples were resting on ice between homogenizer runs), and RNAs were extracted from the collected bacterial cells using TRIzol reagent (Thermo Fisher Scientific, Waltham, MA). RNA was isolated using RNeasy (Qiagen, Germantown, MD) mini spin columns. Sequence libraries were generated using the TruSeq Stranded Total RNA Library Prep kit (Illumina, San Diego, CA) following the manufacturer’s recommendations. The rRNAs were removed by the Ribo-zero Kit (Illumina) to enrich mRNA, using 13 cycles of PCR amplification of the final library. Amplified libraries were purified using AMPure beads (Beckman Coulter, Brea, CA), quantified by Qubit (Thermo Fisher Scientific, Waltham, MA) and qPCR, and visualized in an Agilent Bioanalyzer (Santa Clara, CA). Pooled libraries were sequenced as paired-end 50-bp reads using an Illumina NovaSeq instrument.

Reads were initially trimmed using Trimmomatic version 0.39 (54) to remove adaptors as well as leading or trailing bases with a quality score less than 3, filtering reads with minimum length 36. Reads were mapped to reference strain USA300 FPR3757 (RefSeq identifier GCF_000013465.1) using Bowtie2 version 2.2.5 (55). Using gene annotations from the same assembly, reads mapped to each gene were counted with featureCounts version 2.0.1 (56), producing a counts matrix. Additional analysis was performed in R (R Core Team 2021) using the package DESeq2 version 1.32 (57).

Normalization to account for inter-sample library size variation was performed using the built-in normalization function of DESeq2. All RNA-seq heatmaps were colored according to row (gene) z-scores of DESeq2 normalized counts. For differential expression testing, the Wald test was used with a log-2 fold-change threshold of 0.5 and an FDR of 0.1. For simple pairwise comparisons (e.g., the effect of strain under control conditions), datasets were split so that analysis was performed independently for strains used in the comparison. To determine the interaction between strain and condition variables, all samples were included with the experimental design (formula “expression ∼ condition + strain + condition:strain`”`, where condition:strain is the interaction between variables).

### Metabolic flux prediction

The SPOT (Simplified Pearson cOrrelation with Transcriptomic data) computational method (58) was used to analyze the difference in intracellular metabolic fluxes between wild-type LAC and *agr::tetM* mutant grown in TSB to late exponential phase (OD_600_∼4). SPOT is similar to the E-Flux2 method described previously (59), but a recent validation study (60) shows that SPOT generally outperforms E-Flux2. SPOT infers metabolic flux distribution by integrating transcriptomic data in a genome-scale metabolic model of *S*. *aureus* (61) that was adapted for use with strain LAC. For a list of the metabolic reactions ranked by unit of flux per 100 units of glucose uptake flux, see Supplementary file 3.

### Real-time qRT-PCR assays

Briefly, RNA was purified as described above from late exponential (OD_600_∼4.0) cells, cDNAs were synthesized using Maxima First Strand cDNA Synthesis Kit (Thermo Fisher Scientific, Waltham, MA), and real-time reverse transcription quantitative PCR (qRT-PCR) was performed using QuantiNova™ SYBR Green PCR Kit (Qiagen, Hilden, Germany). Primers were synthesized by IDT Inc. (Coralville, IA). Three independent biological samples were run in triplicate and *rpoB* was used to normalize gene expression. 2^−ΔΔCt^ method was used to calculate the relative fold gene expression (62).

### Peritoneal infection of mice

C57BL/6 mice and C57BL/6 *Cybb*^-/-^ (also known as gp91phox/nox2) were purchased from the Jackson Laboratory and bred onsite to generate animals for experimentation. Age and gender-matched, 8–10-week-old mice were used. *S. aureus* strains harboring *RNAIII* or *agrBD* deletion in the NCTC 8325 background were grown overnight in TSB (37°C, 180 rpm) separately or mixed at a 1:1 ratio. Overnight cultures were diluted (OD_600_∼0.05) into fresh TSB medium (subcultured separately for the cultures mixed overnight (‘primed’) or mixed 1:1 for *RNAIII* or *agrBD* mutant single cultures (‘unprimed’) and grown at 37°C to early exponential phase (OD_600_∼0.5)). Bacteria were washed one time by centrifugation with PBS and adjusted to 10^9^ CFU/mL. Twenty C57BL/6 WT mice and 17 *Cybb*^-/-^ mice were injected intraperitoneally with 100 μL of either ‘primed’ or ‘unprimed’ inoculum. After 2 h, internal organs, peritoneal lavage and blood were collected. The organs were homogenized in sterile PBS and serial dilutions were plated for viable counts on TS agar. Collected blood was lysed with saponin and plated for viable counts on TSA plates. Peritoneal lavage fluid was serially diluted and plated for viable counts. All animal studies were performed as per an NYU Grossman School of Medicine Institutional Animal Care and Use Committee (IACUC) approved protocol to the Shopsin Lab.

### Statistical analysis

Prism software (GraphPad, Inc.) was used to perform statistical analyses. Statistical significance was determined using the Student’s *t* test, Mann–Whitney *U* test, one-way analysis of variance (ANOVA), or the Kruskal-Wallis test, depending on the data type. Statistical significance was considered to be represented by *P* values of <0.05.

### Data availability

The RNA-seq data are available through the NCBI GEO repository using the accession number GSE207045.

## Acknowledgments

We thank Andrew Darwin for critical comments on the manuscript. This work was supported by NIH National Institute of Allergy and Infectious Diseases grants R01AI137336 (B.S., I.Y., and V.J.T.); R01AI140754 (B.S. and V.J.T.); R01AI150893 and R01AI038446 (J.N.W.); R01AI149350 (V.J.T.); K08AI163457 (R.J.U.), and R21AI153646 (D.P.); New Jersey Health Foundation PC 142-22 and New Jersey Commission on Cancer Research COCR22RBG005 grants (D. P.); and funds from the NYU Langone Health Antimicrobial-Resistant Pathogens Program (B.S., A.P., and V.J.T.). The NYU Langone Health Genome Technology Center, and the Cytometry and Cell Sorting Laboratory are shared resources that are partially supported by the Cancer Center Support Grant P30CA016087 at the Laura and Isaac Perlmutter Cancer Center.

## Author Contributions

M.P., Conceptualization, Data curation, Formal analysis, Investigation, Visualization, Methodology, Writing - original draft and review and editing

A.I.P, Investigation, Writing - review and editing

G.P, Data curation, Formal analysis, Visualization, Methodology, Writing - review and editing

A.P., Data curation, Formal analysis, Methodology, Writing - review and editing

J.K., Investigation, Visualization, Writing - review and editing

A.D., Methodology, Investigation, Writing - review and editing

E.Z., Investigation, Writing - review and editing

R.J.U., Investigation, Writing - review and editing

T.K.K., Investigation, Writing - review and editing

C.Z., Investigation, Writing - review and editing

A.F.H., Conceptualization, Writing - review and editing

J.S., Investigation, Writing - review and editing

G.A.S., Resources, Writing - review and editing

J.K., Investigation, Writing - review and editing

J.C., Resources, Investigation, Writing – review and editing

A.R.R., Conceptualization, Investigation, Writing - review and editing

J.N.W., Resources, Writing - review and editing

C.R.N., Investigation, Formal analysis, Visualization, Writing - review and editing

D.S.L., Formal analysis, Visualization, Writing – review and editing

D.P., Supervision, Writing – review and editing

A.P., Supervision, Writing – review and editing

X.Z., Conceptualization, Writing – review and editing

K.D., Conceptualization, Writing – original draft and review and editing

I.Y., Supervision, Funding acquisition

V.J.T., Resources, Supervision, Funding acquisition

B.S., Conceptualization, Resources, Supervision, Funding acquisition, Project Administration, Writing – original draft and review and editing

---

**Figure 1—figure supplement 1.**
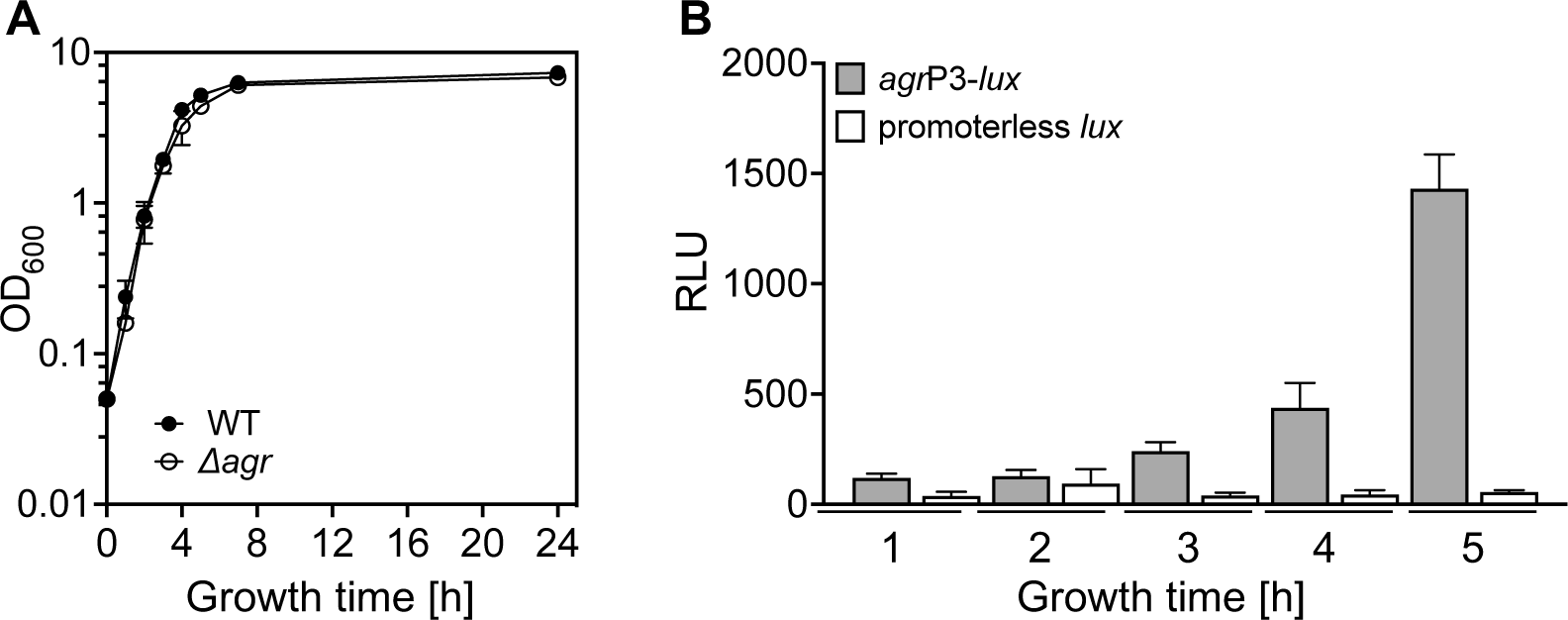
Correlation of growth phase and *agr* expression. (A) Growth curves. Overnight cultures of *S. aureus* LAC wild-type (WT, BS819) or *Δagr* mutant (BS1348) were diluted (OD_600_∼0.05) in fresh TSB medium and growth was monitored by measuring the optical density at 600 nm (OD_600_). (B) Tests of *agr*P3 promoter activity. *S. aureus* LAC wild-type (WT, BS819) containing *agr*P3*-lux* (SaPI1 *att*C::*agr*P3-*lux*; strain BS1222) or control containing a promoterless *lux* gene within the *att*C site (SaPI1 *att*C::pGYLux, strain BS999) grown as in (A) for the indicated times. *agr*P3 activity (relative luminescence units [RLU]) was assayed at the indicated times (see Materials and Methods). Data represent the means ± S.D. from biological replicates (*n* = 3).

**Figure 1—figure supplement 2.**
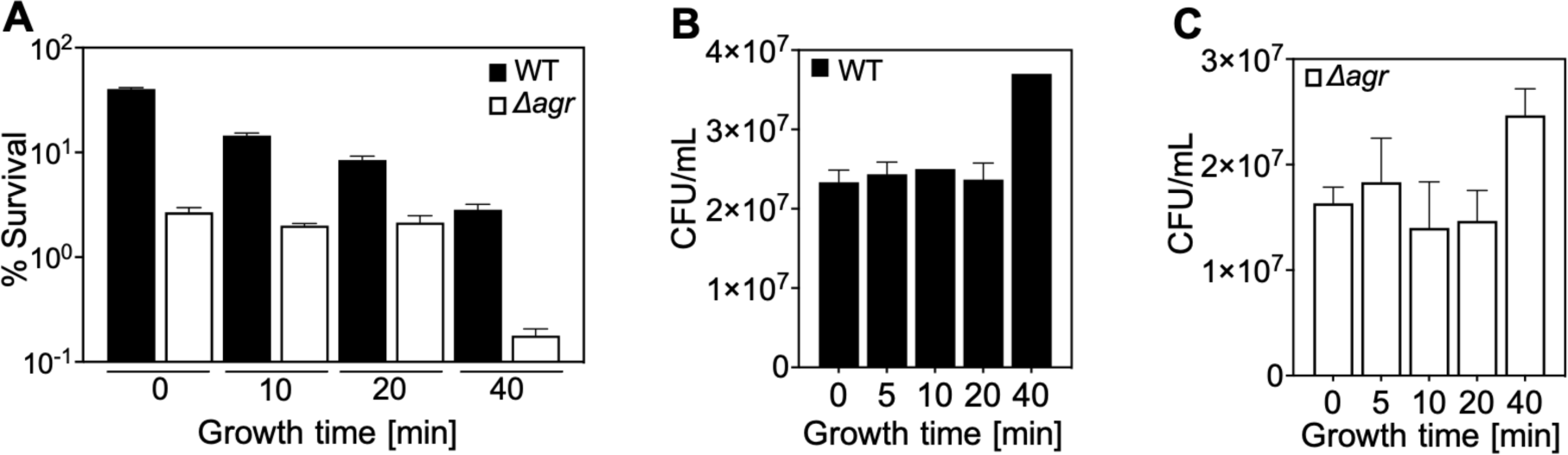
Correlation of lag-time and *agr*-mediated protection from H_2_O_2_-mediated killing. Overnight cultures of *S. aureus* LAC wild-type (WT, BS819) and *Δagr* mutant (BS1348), grown for the indicated times following dilution to fresh medium, were treated with H_2_O_2_ (20 mM for 60 min) (Fig. S1A). Data represent the means ± S.D. from biological replicates (*n* = 3). Survival of Δ*agr* mutant cells was unchanged up to the 40 min time point, and then it dropped sharply. The sharp drop coincided with the time to first division (i.e., the lag time), as evidenced by an increase in colony-forming units (CFUs) at the 40 min time point in the absence of treatment (Fig. S1B and C). In contrast to results with the Δ*agr* strain, survival of the wild-type strain gradually decreased throughout the experiment (Fig. S1A). Increased lag-time is associated with tolerance to lethal stress owing to a delay in growth when switched to a new environment (15). Thus, our observations suggest that a subpopulation of Δ*agr* mutant cells remains longer in a dormant state, decreasing the lethality of H_2_O_2_. The differential effect of the lag time on the wild-type and Δ*agr* mutant cultures was absent during exponential growth (40 min). These results suggest that *agr* contributes to at least two forms of protection from H_2_O_2_-mediated killing: tolerance by a transient lag state and tolerance during growth phase. To focus on the latter form, assays involving cultures after overnight growth were grown for ∼65 min (OD_600_∼0.15).

**Figure 1—figure supplement 3.**
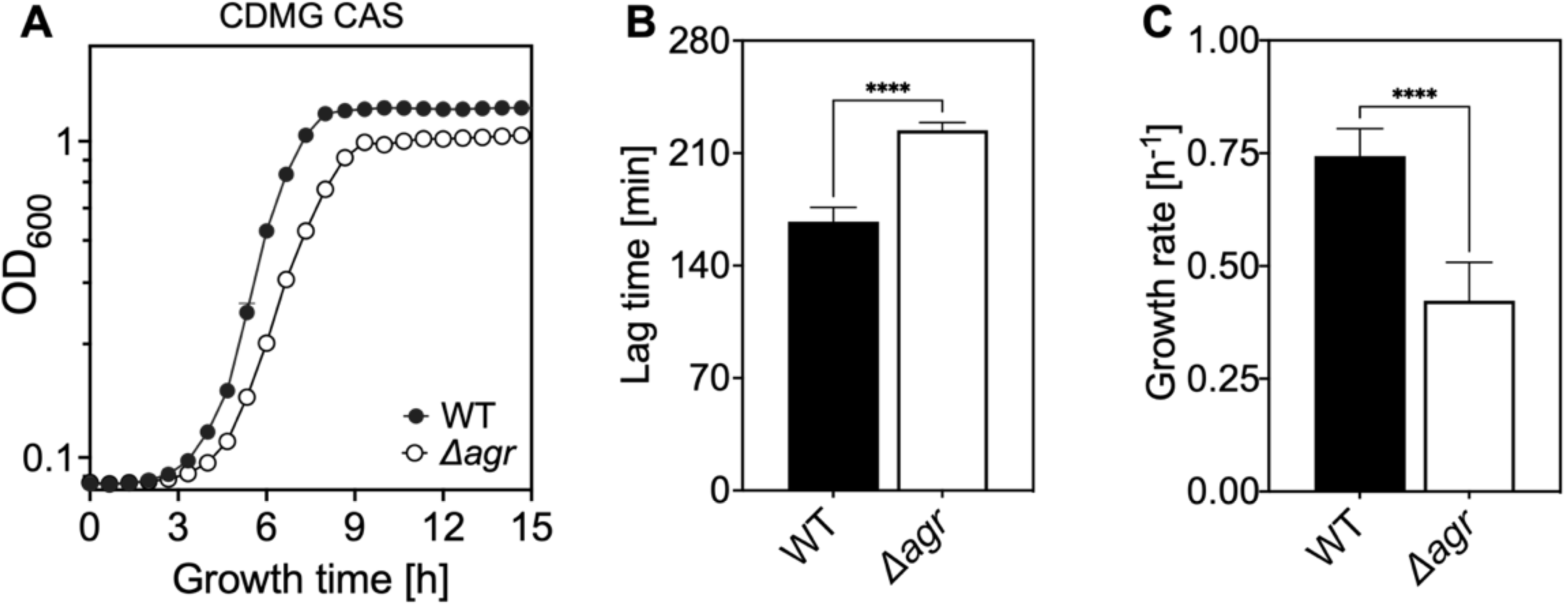
Extended lag phase and decreased growth rate and yield of an Δ*agr* mutant. (A) Growth curves. *S. aureus* LAC wild-type (WT, BS819) and *Δagr* mutant (BS1348) cultures were grown in chemically-defined medium supplemented with 0.5% Casamino acids and 14 mM glucose (CDMG CAS) for the indicated times following 1,000-fold dilution of overnight cultures grown in TSB. Growth of diluted cultures was monitored for 15 hours every 40 minutes by measuring the OD_600_ using an Agilent LogPhase 600 Microbiology Reader (Santa Clara, CA). (B) Lag times. Data in panel A were used to determine lag times by extrapolation of the linear portion of the growth curve. Growth rates (µ, h^−1^) calculated from five biological replicates are displayed in panel (B). Data are means ± S.D. Statistical significance was calculated with Student’s two-tailed *t* test (****, *P* ≤ 0.0001).

**Figure 1—figure supplement 4.**
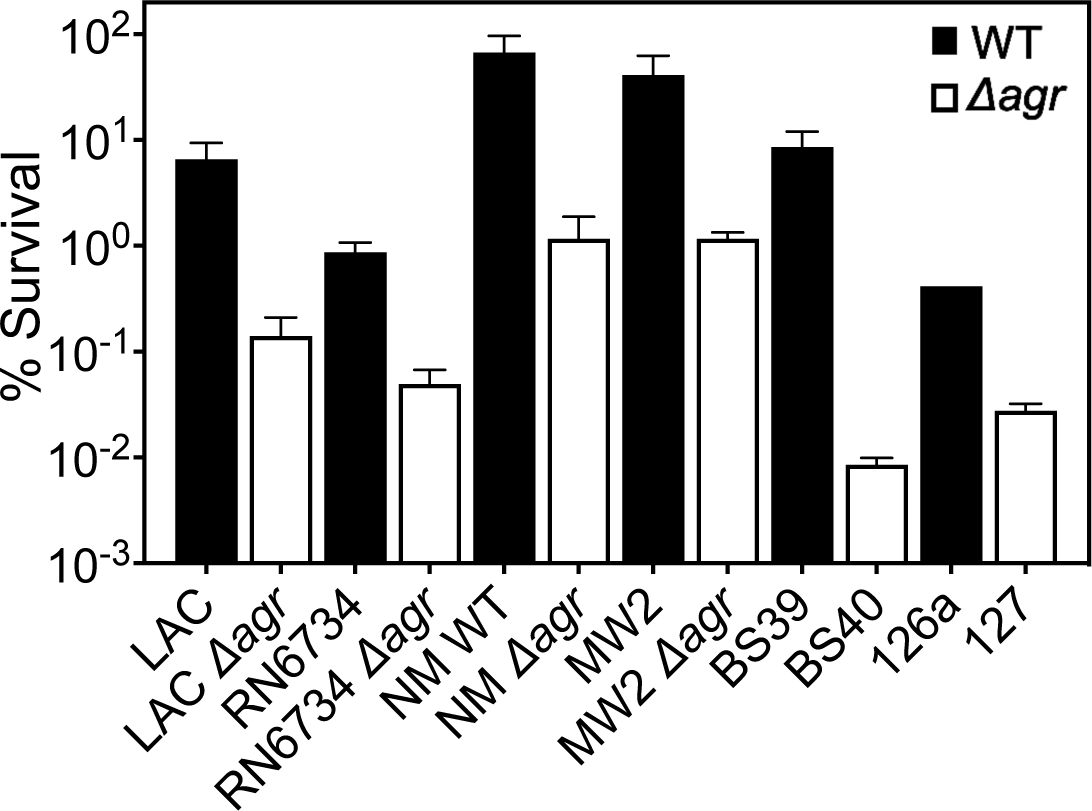
*Agr*-mediated protection from H_2_O_2_-mediated killing among diverse *S. aureus* strains. Laboratory strains LAC, RN6734, Newman (NM, BS12), MW2 (BS450), and clinical isolates BS39 and 126a with *agr* deficient mutant derivatives were compared for survival following treatment with 20 mM of H_2_O_2_ for 60 min. In this experiment overnight cultures were diluted in TSB and grown to early log phase (OD_600_∼0.15). Percent survival was determined relative to samples taken at the time of peroxide treatment. Some mutants were created by transduction of marker-disrupted alleles (LAC, RN6734, Newman, MW2) while others were naturally occurring (BS40, 127) (see Table 1). Data represent the means ± S.D. from biological replicates (*n* = 3). The data show that peroxide lethality varies among strains, but in each case deletion of *agr* increases killing.

**Figure 2—figure supplement 1.**
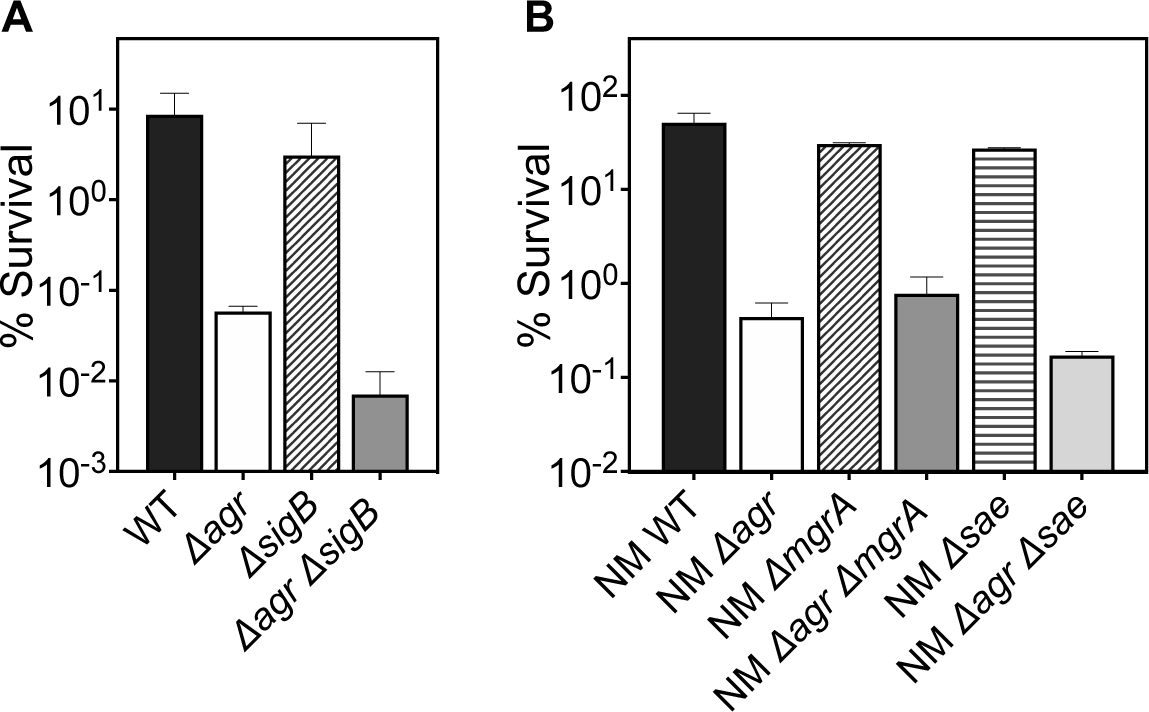
Deficiency of downstream global regulators does not differentially affect *agr*-mediated protection from H_2_O_2_-mediated cell death. The effect of (A) *sigB*, (B) *mgrA* and *sae* on survival in the presence or absence of *agr* during treatment with H_2_O_2_ was measured. Cells were grown to early log phase (OD_600_∼0.15) and treated with 20 mM of H_2_O_2_ for 60 min. Data represent the means ± S.D. from biological replicates (*n* = 3). Bacterial strains were BS819 (LAC) and BS12 (Newman, NM) for WT and BS1348 (LAC) and BS13 (NM) for the *agr*, and BS BS1435-36, BS1280, BS1282 and BS1246, BS1518 for *sigB*, *sae* and *mgrA* mutants, respectively. The genes tested were either part of known two-component systems or SarA protein-family regulatory circuits involved in virulence gene expression. They are all downstream/epistatic to *agr*-RNAIII (reviewed in (17)). Mutations in *sigB*, *mgrA*, and *sae* showed little or no effect with respect to the protective *agr*-mediated phenotype. These results support the idea that *rot* is the primary regulator pathway that protects the wild-type from H_2_O_2_-mediated killing.

**Figure 4—figure supplement 1.**
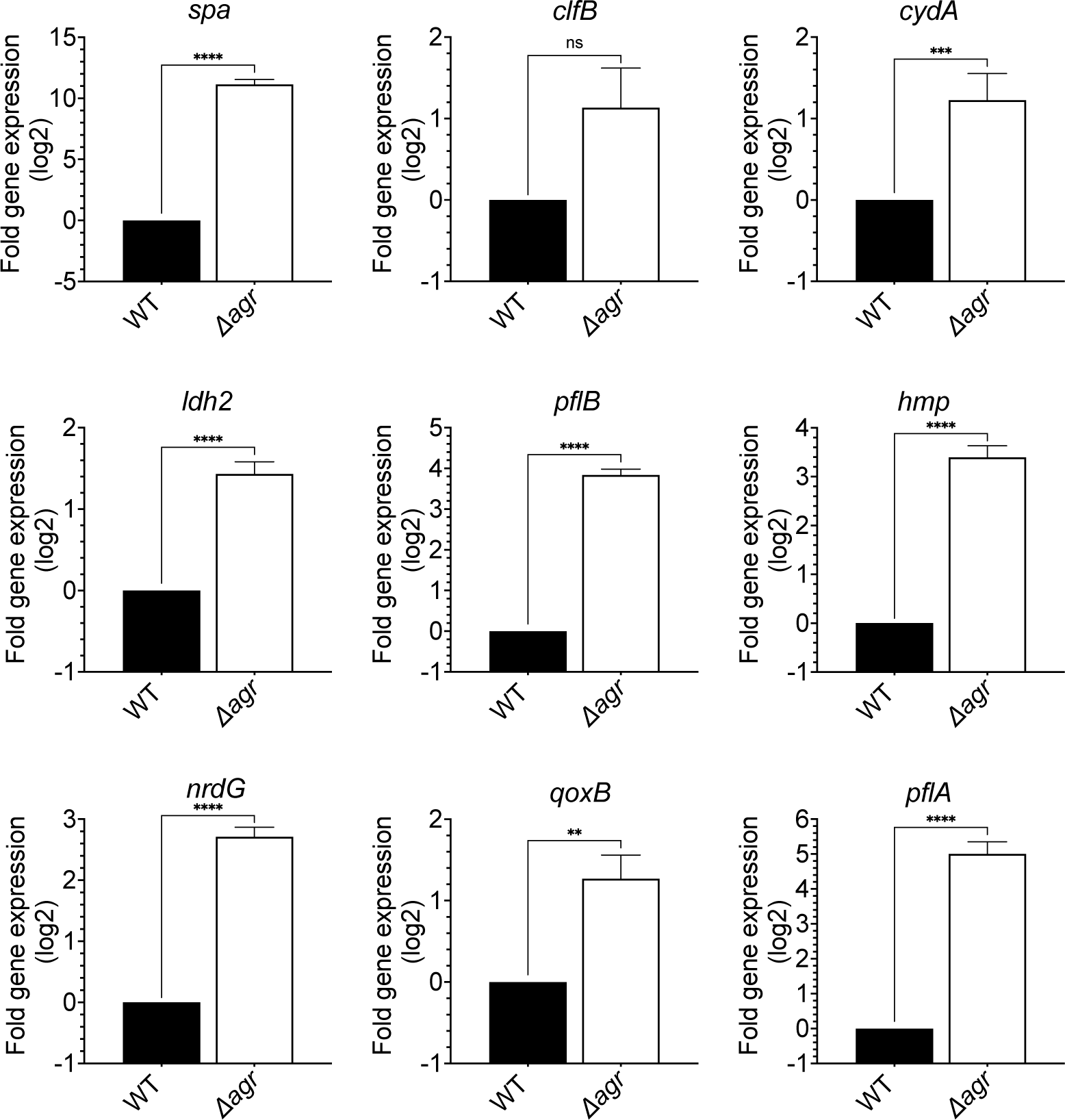
Induction of expression of selected fermentive/anaerobic genes stimulated by deletion of *agr*. Total cellular RNA was extracted from late exponential phase cultures (OD_600_∼4.0) of wild-type (WT, BS819) or Δ*agr* mutant (BS1348), followed by reverse transcription and PCR amplification of the indicated genes from Fig. 5A, using *rpoB* as an internal standard. mRNA levels were normalized to those of each gene with an untreated wild-type control. Data represent the mean ± SEM of three independent experiments. Student’s t test was used to determine statistical differences between samples (\*\**P* < 0.01; \*\*\**P* < 0.001; \*\*\*\**P* < 0.0001). In each case the *agr* deletion increased expression, indicating elevated metabolism.

**Figure 5—figure supplement 1.**
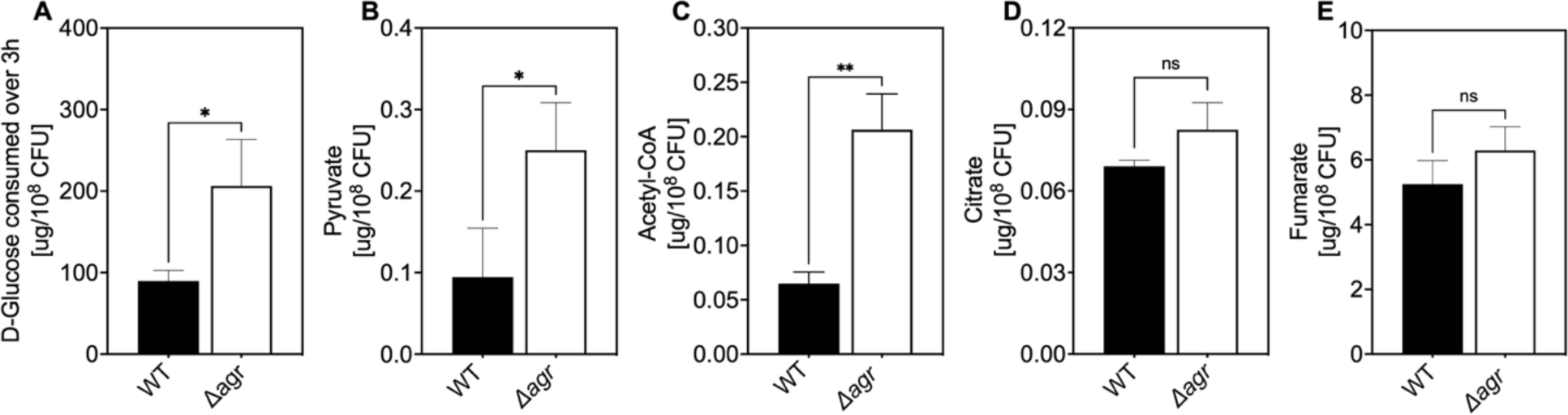
Association of *agr* deficiency with glucose consumption and intracellular levels of pyruvate, acetyl-CoA, and TCA-cycle metabolites. (A) Extracellular glucose levels. D-glucose levels per expressed as expressed as µg/10^8^ cells of *S. aureus* LAC wild-type (WT, BS819) or Δ*agr* mutant (BS1348) after growth of cultures in TSB medium to exponential phase (3 h). (B-E) Intracellular levels of pyruvate, acetyl-CoA, and TCA-cycle metabolites citrate and fumarate. Levels of the indicated metabolite expressed as expressed as µg/10^8^ cells of *S. aureus* LAC wild-type (WT, BS819) or Δ*agr* mutant (BS1348) after growth in TSB medium to late-exponential phase (OD_600_∼4.0). For all panels, data points are the mean value ± SD (*n* = 3). \**P* < 0.05; \*\*\*\**P* < 0.0001, by Student’s two-tailed *t* test.

**Figure 7—figure supplement 1.**
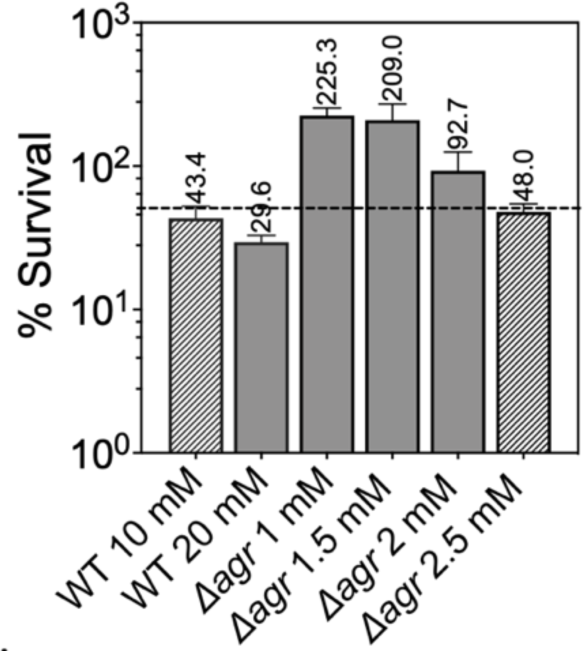
Normalization of the lethal concentration of H_2_O_2_ with wild-type and Δ*agr* strains. Overnight cultures of were diluted into fresh TSB medium and grown to early log phase (OD_600_ = 0.15 to achieve sufficient CFU for RNA-seq). These cultures were treated with the indicated concentrations of H_2_O_2_ for 30 min prior to measurement of survival by plating. Data represent the means ± S.D. from biological replicates (*n* = 3). Bacterial strains were BS819 for WT and BS1348 for the *agr* mutant. To focus RNA-seq analysis on lethal rather than cell death responses, we sought to reduce H_2_O_2_ concentrations and thereby lethality to achieve ∼50% (dotted line) cell survival, normalized to wild-type and Δ*agr* mutant strains. Survival of the Δ*agr* mutant with H_2_O_2_ for 30 min at a concentration of 2.5 mM closely approximated 50% survival of the wild-type with 10 mM H_2_O_2,_ providing a basis for choice of concentrations and treatment time for RNA-seq analysis.

**Figure 8—figure supplement 1.**
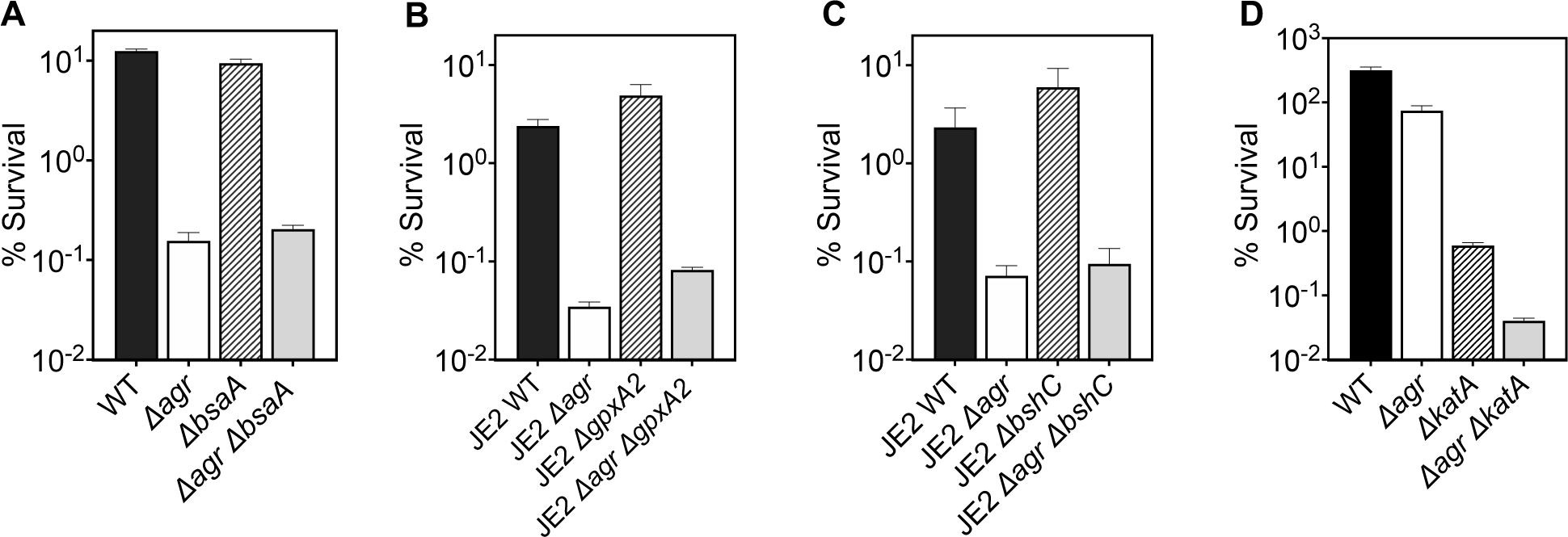
Deficiency in ROS detoxification genes (*katA, bsaA1/gpxA1, bsaA2/gpxA2*, and bacilliothiol (BSH) have no effect on *agr*-mediated protection from H_2_O_2_-mediated cell death. Effect of (A) *bsaA* (B) *gpxA2*, (C) *bshC* and (D) *katA* on survival during treatment with H_2_O_2_. Cells were grown to early log phase (OD_600_∼0.15) and treated with 20 mM of H_2_O_2_ for 60 min for A-C or with 2 mM of H_2_O_2_ for 60 min for D. Data represent the means ± S.D. from biological replicates (*n* = 3). Bacterial strains were BS819 (LAC) and BS867 (LAC) for WT, and BS1348 (LAC), BS1010 (JE2), BS1490-91, BS1522-23, BS1527-28 and BS1488-89 for the *agr*, *bsaA*, *gpxA2*, *bshC*, and *katA* mutants, respectively. Our data with superoxide dismutases (*sodA*) and the peroxiredoxin *ahpC* (Fig. 7) suggest that homeostatic detoxification pathways contribute to *agr*-mediated phenotypes with respect to lethal H_2_O_2_ stress. Mutations in additional genes involved in H_2_O_2_ detoxification that included catalase, (*katA*), two thiol-dependent peroxidases (*gpxA1* and *gpxA2*), and the low-molecular-weight thiol bacillithiol (*bshC*) showed no differential effect with respect to *agr*-mediated phenotypes. Notably, *gpxA1,* which is also known as *bsaA1,* was essential for the oxidation-sensing ability of AgrA to confer resistance to H_2_O_2_-mediated growth inhibition (7). The Δ*katA* mutation was hyperlethal with the wild-type and Δ*agr* mutant, even when otherwise sub-inhibitory concentrations of H_2_O_2_ were used. Collectively, the data support the idea that *agr*-mediated phenotypes are detoxification pathway-specific.

**Figure 8—figure supplement 2.**
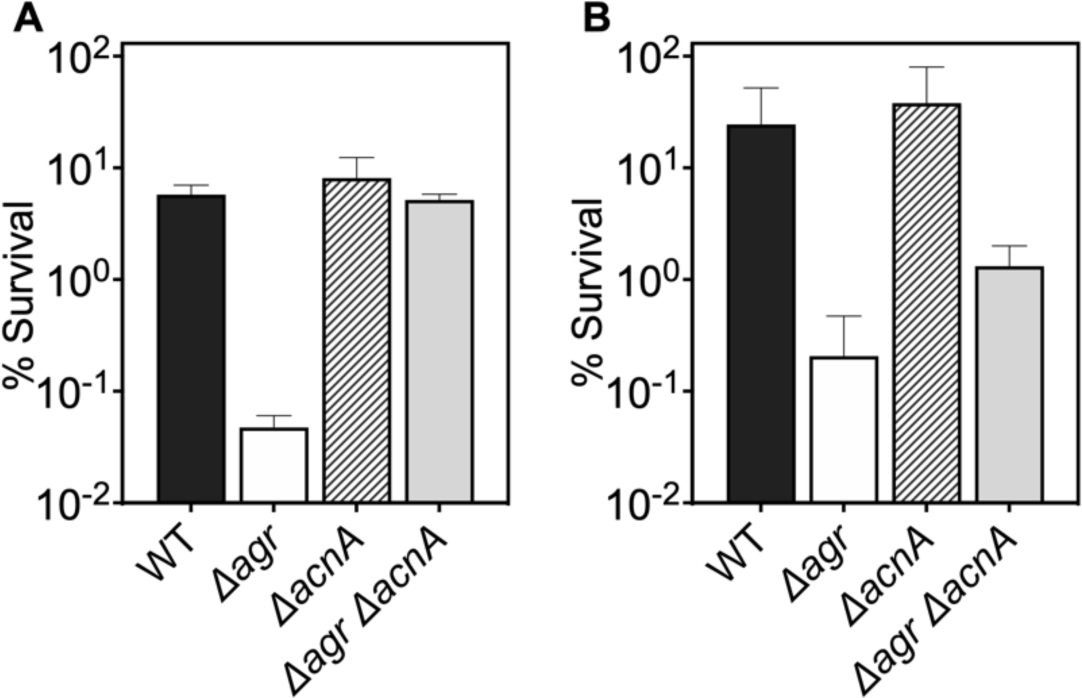
Deficiency in TCA cycle gene *acnA* reverses the effect of an *agr* deficiency with respect to subsequent challenge with H_2_O_2_. (A-B) Effect of *acnA* on survival during treatment with H_2_O_2_. Cells were grown to (A) late (OD_600_∼4.0) or (B) early (OD_600_∼0.15) log phase and treated with 20 mM of H_2_O_2_ for 60 min. Bacterial strains were *S. aureus* LAC wild-type (WT, BS819), Δ*agr* mutant (BS1348), *acnA::bursa* (BS1744), and *acnA::bursa-Δagr* double mutant (BS1745). Data represent the means ± S.D. from biological replicates (*n* = 3).

**Figure 8—figure supplement 3.**
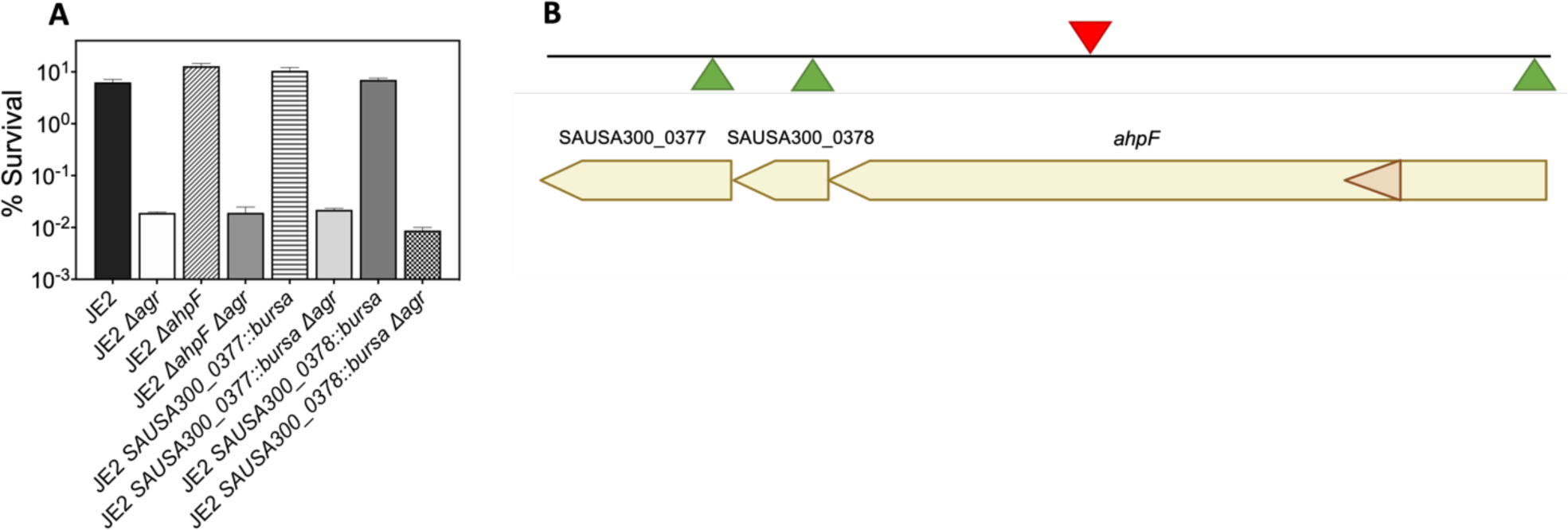
Effects of transposon insertion in *ahpC* unexplained by polarity of transposon insertion. (A) Cultures of *S. aureus* wild-type, *Δagr* mutant, and various double mutants were treated with H_2_O_2_ (20 mM for 60 min) prior to measurement of survival. For strain descriptions, see *SI* Appendix, Table S2. (B) *ahpC* locus map showing the three ORFs located downstream of *ahpC*. The location of four *Bursa aurealis* insertions (NE911, NE1571, NE537, NE725), obtained from The Nebraska Transposon Mutant Library (NTML) (51) used in this study are indicated by triangles. Green triangles, plus-strand insertion; red triangle, minus-strand insertion. Data represent the means ± S.D. from biological replicates (*n* = 3). Bacterial strains were BS435 for WT and BS1010, BS1494, BS1504, BS1495, BS1501, BS1496 and BS1506 for the *agr*, *ahpF*, SAUSA300_0377, and SAUSA300_0378 mutants, respectively. Since the *Bursa aurealis* (bursa) transposon insertion in *ahpC* was upstream of several open reading frames (ORFs) in the *ahp*C-F operon, polarity could complicate interpretation of the results. We therefore analyzed the effects of the three bursa mutants in strain JE2 downstream genes: *ahpF*, SAUSA300_0378, SAUSA300_0377. We found that polar effects on downstream elements could not explain the properties of *ahpC*::*bursa*. Thus, *ahpC*::*bursa* could provide insights into the role of *ahpC* in *agr*-mediated phenotypes.

**Figure 9—figure supplement 1.**
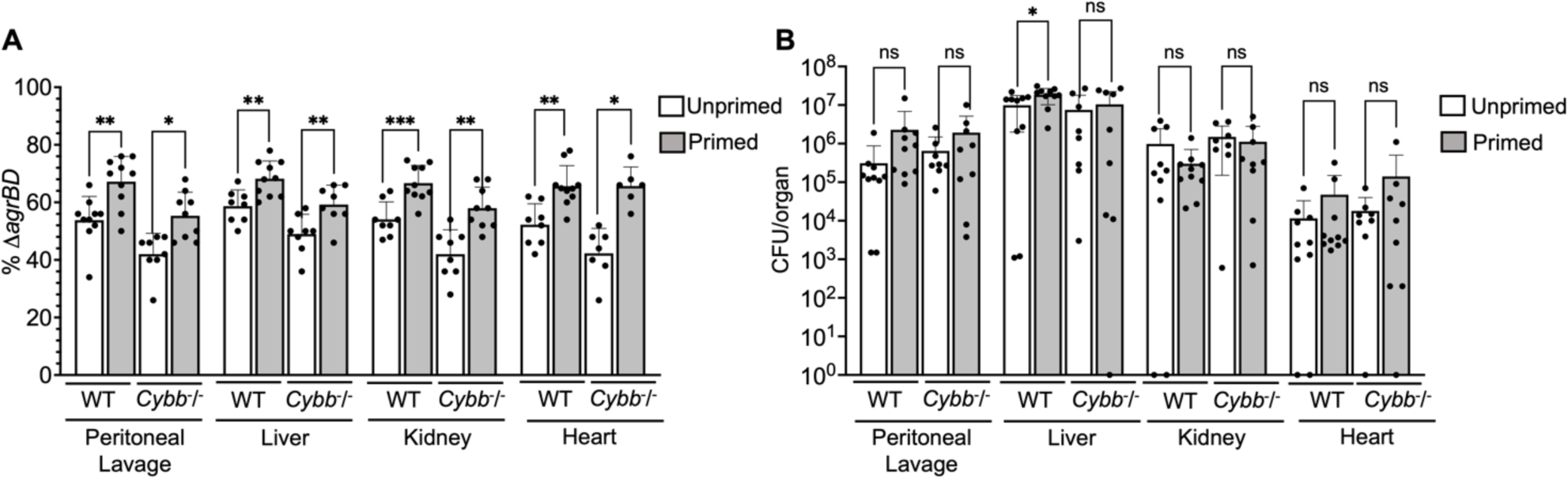
Long-lived protection by *agr* increases peritoneal fitness and dissemination to liver, kidney, and heart in both C57BL/6 mice and C57BL/6 *Cybb^-/-^* (*gp91phox/nox2*) mice. (A) Percentage of 11*agrBD* or (B) CFU of *S. aureus* RN6734 *ΔrnaIII* (GAW183) and *ΔagrBD* mutant (GAW130) cells in the indicated organ 1 h post intraperitoneal infection of wild-type (WT) C57BL/6 mice or phagocyte NADPH oxidase deficient (*Cybb^-/-^*) mice (see Fig. 9 for data with lung and spleen). Wild-type and mutant strains were grown separately and mixed in a 1:1 ratio either before or after overnight growth, as for Fig. 4. We called wild-type and mutant populations that were mixed prior to or after overnight growth; they were termed “primed” and “unprimed”, respectively. Both primed and unprimed mixtures were subsequently diluted, grown to early log phase (OD_600_∼0.15), and used as inoculum for intraperitoneal infection with 1 x 10^8^ CFU (*n* = 2 groups of 10 mice each). After 1h, the peritoneum was lavaged and the heart, kidneys, liver, lungs and spleen (Fig. 9) were harvested and homogenized. Samples were then diluted and plated to enumerate viable bacteria. Output ratios and total and mutant CFU from tissue homogenates were determined as for Fig 4E and 4H. A Mann-Whitney test (9A) or Student’s two-tailed *t* test (9B) were used to determine the statistical significance of the difference between primed and unprimed cultures. Error bars indicate standard deviation (\*\**P* < 0.05; \*\*\*\**P* < 0.0001). Long-lived *agr*-mediated functions increased *S. aureus* pathogenesis in both wild-type and mutant mice, indicating a role for long-lived *agr*-mediated functions in pathogenesis other than protection from ROS. Additionally, long-lived *agr*-mediated protection against ROS enhances fitness in lung and spleen (Fig. 9), but it is dispensable for full virulence in other organs; protection is tissue-specific.

## References

1. Spaan AN, Surewaard BG, Nijland R, van Strijp JA. 2013. Neutrophils versus *Staphylococcus aureus*: a biological tug of war. Annu Rev Microbiol 67:629–650.

2. Drlica K, Zhao X. 2021. Bacterial death from treatment with fluoroquinolones and other lethal stressors. Expert review of anti-infective therapy 19:601–618.

3. Kohanski MA, Dwyer DJ, Hayete B, Lawrence CA, Collins JJ. 2007. A common mechanism of cellular death induced by bactericidal antibiotics. Cell 130:797–810.

4. Lobritz MA, Belenky P, Porter CB, Gutierrez A, Yang JH, Schwarz EG, Dwyer DJ, Khalil AS, Collins JJ. 2015. Antibiotic efficacy is linked to bacterial cellular respiration. Proc Natl Acad Sci U S A 112:8173–8180.

5. Shatalin K, Nuthanakanti A, Kaushik A, Shishov D, Peselis A, Shamovsky I, Pani B, Lechpammer M, Vasilyev N, Shatalina E, Rebatchouk D, Mironov A, Fedichev P, Serganov A, Nudler E. 2021. Inhibitors of bacterial H(2)S biogenesis targeting antibiotic resistance and tolerance. Science 372:1169–1175.

6. Shee S, Singh S, Tripathi A, Thakur C, Kumar TA, Das M, Yadav V, Kohli S, Rajmani RS, Chandra N, Chakrapani H, Drlica K, Singh A. 2022. Moxifloxacin-mediated killing of *Mycobacterium tuberculosis* involves respiratory downshift, reductive stress, and Accumulation of reactive oxygen species. Antimicrob Agents Chemother 66:e0059222.

7. Sun F, Liang H, Kong X, Xie S, Cho H, Deng X, Ji Q, Zhang H, Alvarez S, Hicks LM, Bae T, Luo C, Jiang H, He C. 2012. Quorum-sensing *agr* mediates bacterial oxidation response via an intramolecular disulfide redox switch in the response regulator AgrA. Proc Natl Acad Sci U S A 109:9095–9100.

8. Khan BA, Yeh AJ, Cheung GY, Otto M. 2015. Investigational therapies targeting quorum-sensing for the treatment of *Staphylococcus aureus* infections. Expert Opin Investig Drugs 24:689–704.

9. Novick RP. 2003. Autoinduction and signal transduction in the regulation of staphylococcal virulence. Mol Microbiol 48:1429–1449.

10. Novick RP, Geisinger E. 2008. Quorum sensing in staphylococci. Annu Rev Genet 42:541–564.

11. Novick RP, Projan SJ, Kornblum J, Ross HF, Ji G, Kreiswirth B, Vandenesch F, Moghazeh S. 1995. The agr P2 operon: an autocatalytic sensory transduction system in *Staphylococcus aureus*. Mol Gen Genet 248:446–458.

12. Kumar K, Chen J, Drlica K, Shopsin B. 2017. Tuning of the lethal response to multiple stressors with a single-site mutation during clinical infection by *Staphylococcus aureus*. mBio 8.

13. Geisinger E, Chen J, Novick RP. 2012. Allele-dependent differences in quorum-sensing dynamics result in variant expression of virulence genes in *Staphylococcus aureus*. J Bacteriol 194:2854–2864.

14. Conlon BP, Rowe SE, Gandt AB, Nuxoll AS, Donegan NP, Zalis EA, Clair G, Adkins JN, Cheung AL, Lewis K. 2016. Persister formation in *Staphylococcus aureus* is associated with ATP depletion. Nat Microbiol 1.

15. Fridman O, Goldberg A, Ronin I, Shoresh N, Balaban NQ. 2014. Optimization of lag time underlies antibiotic tolerance in evolved bacterial populations. Nature 513:418–421.

16. Geisinger E, Adhikari RP, Jin R, Ross HF, Novick RP. 2006. Inhibition of *rot* translation by *RNAIII*, a key feature of *agr* function. Mol Microbiol 61:1038–1048.

17. Bronesky D, Wu Z, Marzi S, Walter P, Geissmann T, Moreau K, Vandenesch F, Caldelari I, Romby P. 2016. *Staphylococcus aureus RNAIII* and its regulon link quorum sensing, stress responses, metabolic adaptation, and regulation of virulence gene expression. Annu Rev Microbiol 70:299–316.

18. Fuchs S, Pane-Farre J, Kohler C, Hecker M, Engelmann S. 2007. Anaerobic gene expression in *Staphylococcus aureus*. J Bacteriol 189:4275–4289.

19. Pagels M, Fuchs S, Pane-Farre J, Kohler C, Menschner L, Hecker M, McNamarra PJ, Bauer MC, von Wachenfeldt C, Liebeke M, Lalk M, Sander G, von Eiff C, Proctor RA, Engelmann S. 2010. Redox sensing by a Rex-family repressor is involved in the regulation of anaerobic gene expression in *Staphylococcus aureus*. Mol Microbiol 76:1142–1161.

20. Somerville GA, Beres SB, Fitzgerald JR, DeLeo FR, Cole RL, Hoff JS, Musser JM. 2002. *In vitro* serial passage of *Staphylococcus aureus*: changes in physiology, virulence factor production, and *agr* nucleotide sequence. J Bacteriol 184:1430–1437.

21. Sadykov MR, Thomas VC, Marshall DD, Wenstrom CJ, Moormeier DE, Widhelm TJ, Nuxoll AS, Powers R, Bayles KW. 2013. Inactivation of the Pta-AckA pathway causes cell death in *Staphylococcus aureus*. J Bacteriol 195:3035–3044.

22. Somerville GA, Chaussee MS, Morgan CI, Fitzgerald JR, Dorward DW, Reitzer LJ, Musser JM. 2002. *Staphylococcus aureus* aconitase inactivation unexpectedly inhibits post-exponential-phase growth and enhances stationary-phase survival. Infect Immun 70:6373–6382.

23. Posada AC, Kolar SL, Dusi RG, Francois P, Roberts AA, Hamilton CJ, Liu GY, Cheung A. 2014. Importance of bacillithiol in the oxidative stress response of *Staphylococcus aureus*. Infect Immun 82:316–332.

24. Rowe SE, Wagner NJ, Li L, Beam JE, Wilkinson AD, Radlinski LC, Zhang Q, Miao EA, Conlon BP. 2020. Reactive oxygen species induce antibiotic tolerance during systemic *Staphylococcus aureus* infection. Nat Microbiol 5:282–290.

25. Clements MO, Watson SP, Foster SJ. 1999. Characterization of the major superoxide dismutase of *Staphylococcus aureus* and its role in starvation survival, stress resistance, and pathogenicity. J Bacteriol 181:3898–3903.

26. Gaupp R, Ledala N, Somerville GA. 2012. Staphylococcal response to oxidative stress. Frontiers in cellular and infection microbiology 2:33.

27. Imlay JA. 2008. Cellular defenses against superoxide and hydrogen peroxide. Annu Rev Biochem 77:755–776.

28. Cosgrove K, Coutts G, Jonsson IM, Tarkowski A, Kokai-Kun JF, Mond JJ, Foster SJ. 2007. Catalase (KatA) and alkyl hydroperoxide reductase (AhpC) have compensatory roles in peroxide stress resistance and are required for survival, persistence, and nasal colonization in *Staphylococcus aureus*. J Bacteriol 189:1025–1035.

29. Seaver LC, Imlay JA. 2001. Alkyl hydroperoxide reductase is the primary scavenger of endogenous hydrogen peroxide in *Escherichia coli*. J Bacteriol 183:7173–7181.

30. Pollock JD, Williams DA, Gifford MA, Li LL, Du X, Fisherman J, Orkin SH, Doerschuk CM, Dinauer MC. 1995. Mouse model of X-linked chronic granulomatous disease, an inherited defect in phagocyte superoxide production. Nature genetics 9:202–209.

31. Yipp BG, Kim JH, Lima R, Zbytnuik LD, Petri B, Swanlund N, Ho M, Szeto VG, Tak T, Koenderman L, Pickkers P, Tool ATJ, Kuijpers TW, van den Berg TK, Looney MR, Krummel MF, Kubes P. 2017. The lung is a host defense niche for immediate neutrophil-mediated vascular protection. Sci Immunol 2.

32. Beavers WN, DuMont AL, Monteith AJ, Maloney KN, Tallman KA, Weiss A, Christian AH, Toste FD, Chang CJ, Porter NA, Torres VJ, Skaar EP. 2021. *Staphylococcus aureus* peptide methionine sulfoxide reductases protect from human whole-blood killing. Infect Immun 89:e0014621.

33. Zeng J, Hong Y, Zhao N, Liu Q, Zhu W, Xiao L, Wang W, Chen M, Hong S, Wu L, Xue Y, Wang D, Niu J, Drlica K, Zhao X. 2022. A broadly applicable, stress-mediated bacterial death pathway regulated by the phosphotransferase system (PTS) and the cAMP-Crp cascade. Proc Natl Acad Sci U S A 119:e2118566119.

34. Morfeldt E, Taylor D, von Gabain A, Arvidson S. 1995. Activation of alpha-toxin translation in *Staphylococcus aureus* by the trans-encoded antisense RNA, *RNAIII*. EMBO J 14:4569–4577.

35. Novick RP, Ross HF, Projan SJ, Kornblum J, Kreiswirth B, Moghazeh S. 1993. Synthesis of staphylococcal virulence factors is controlled by a regulatory RNA molecule. EMBO J 12:3967–3975.

36. Takahashi N, Gruber CC, Yang JH, Liu X, Braff D, Yashaswini CN, Bhubhanil S, Furuta Y, Andreescu S, Collins JJ, Walker GC. 2017. Lethality of MalE-LacZ hybrid protein shares mechanistic attributes with oxidative component of antibiotic lethality. Proc Natl Acad Sci U S A 114:9164–9169.

37. George SE, Hrubesch J, Breuing I, Vetter N, Korn N, Hennemann K, Bleul L, Willmann M, Ebner P, Gotz F, Wolz C. 2019. Oxidative stress drives the selection of quorum sensing mutants in the *Staphylococcus aureus* population. Proc Natl Acad Sci U S A 116:19145–19154.

38. Queck SY, Jameson-Lee M, Villaruz AE, Bach TH, Khan BA, Sturdevant DE, Ricklefs SM, Li M, Otto M. 2008. *RNAIII*-independent target gene control by the *agr* quorum-sensing system: insight into the evolution of virulence regulation in *Staphylococcus aureus*. Mol Cell 32:150–158.

39. Wang R, Braughton KR, Kretschmer D, Bach T-HL, Queck SY, Li M, Kennedy AD, Dorward DW, Klebanoff SJ, Peschel A, DeLeo FR, Otto M. 2007. Identification of novel cytolytic peptides as key virulence determinants for community-associated MRSA. Nat Med 13:1510–1514.

40. Dorsey-Oresto A, Lu T, Mosel M, Wang X, Salz T, Drlica K, Zhao X. 2013. YihE kinase is a central regulator of programmed cell death in bacteria. Cell reports 3:528–537.

41. Wu X, Wang X, Drlica K, Zhao X. 2011. A toxin-antitoxin module in *Bacillus subtilis* can both mitigate and amplify effects of lethal stress. PLoS One 6:e23909.

42. Richardson AR, Libby SJ, Fang FC. 2008. A nitric oxide-inducible lactate dehydrogenase enables *Staphylococcus aureus* to resist innate immunity. Science 319:1672–1676.

43. Vitko NP, Spahich NA, Richardson AR. 2015. Glycolytic dependency of high-level nitric oxide resistance and virulence in *Staphylococcus aureus*. mBio 6.

44. Vilcheze C, Jacobs WR, Jr. 2021. The promises and limitations of N-acetylcysteine as a potentiator of first-line and second-line tuberculosis drugs. Antimicrob Agents Chemother.

45. Cao S, Huseby DL, Brandis G, Hughes D. 2017. Alternative evolutionary pathways for drug-resistant small colony variant mutants in *Staphylococcus aureus*. mBio 8.

46. Gusarov I, Shatalin K, Starodubtseva M, Nudler E. 2009. Endogenous nitric oxide protects bacteria against a wide spectrum of antibiotics. Science 325:1380–1384.

47. Shatalin K, Shatalina E, Mironov A, Nudler E. 2011. H2S: a universal defense against antibiotics in bacteria. Science 334:986–990.

48. Novick RP. 1991. Genetic systems in staphylococci. Methods Enzymol 204:587–636.

49. Hussain M, Hastings JG, White PJ. 1991. A chemically defined medium for slime production by coagulase-negative staphylococci. J Med Microbiol 34:143–147.

50. Grosser MR, Weiss A, Shaw LN, Richardson AR. 2016. Regulatory requirements for *Staphylococcus aureus* nitric oxide resistance. J Bacteriol 198:2043–2055.

51. Fey PD, Endres JL, Yajjala VK, Widhelm TJ, Boissy RJ, Bose JL, Bayles KW. 2013. A genetic resource for rapid and comprehensive phenotype screening of nonessential *Staphylococcus aureus* genes. mBio 4:e00537–00512.

52. Chen J, Yoong P, Ram G, Torres VJ, Novick RP. 2014. Single-copy vectors for integration at the SaPI1 attachment site for *Staphylococcus aureus*. Plasmid 76:1–7.

53. Geisinger E, George EA, Muir TW, Novick RP. 2008. Identification of ligand specificity determinants in AgrC, the *Staphylococcus aureus* quorum-sensing receptor. J Biol Chem 283:8930–8938.

54. Bolger AM, Lohse M, Usadel B. 2014. Trimmomatic: a flexible trimmer for Illumina sequence data. Bioinformatics 30:2114–2120.

55. Langmead B, Salzberg SL. 2012. Fast gapped-read alignment with Bowtie 2. Nat Methods 9:357–359.

56. Liao Y, Smyth GK, Shi W. 2014. featureCounts: an efficient general purpose program for assigning sequence reads to genomic features. Bioinformatics 30:923–930.

57. Love MI, Huber W, Anders S. 2014. Moderated estimation of fold change and dispersion for RNA-seq data with DESeq2. Genome Biol 15:550.

58. Kim MK, Lane A, Kelley JJ, Lun DS. 2016. E-Flux2 and SPOT: Validated methods for inferring intracellular metabolic flux distributions from transcriptomic data. PLoS One 11:e0157101.

59. Balasubramanian D, Ohneck EA, Chapman J, Weiss A, Kim MK, Reyes-Robles T, Zhong J, Shaw LN, Lun DS, Ueberheide B, Shopsin B, Torres VJ. 2016. *Staphylococcus aureus* coordinates leukocidin expression and pathogenesis by sensing metabolic fluxes via RpiRc. mBio 7.

60. Bhadra-Lobo S, Kim MK, Lun DS. 2020. Assessment of transcriptomic constraint-based methods for central carbon flux inference. PLoS One 15:e0238689.

61. Becker SA, Palsson BO. 2005. Genome-scale reconstruction of the metabolic network in *Staphylococcus aureus* N315: an initial draft to the two-dimensional annotation. BMC microbiology 5:8.

62. Livak KJ, Schmittgen TD. 2001. Analysis of relative gene expression data using real-time quantitative PCR and the 2^−ΔΔ*CT*^ Method. Methods 25:402–408.

63. Wright JS, 3rd, Jin R, Novick RP. 2005. Transient interference with staphylococcal quorum sensing blocks abscess formation. Proc Natl Acad Sci U S A 102:1691–1696.

64. Andrade-Linares DR, Lehmann A, Rillig MC. 2016. Microbial stress priming: a meta-analysis. Environ Microbiol 18:1277–1288.

65. Boles BR, Thoendel M, Roth AJ, Horswill AR. 2010. Identification of genes involved in polysaccharide-independent *Staphylococcus aureus* biofilm formation. PLoS One 5:e10146.

66. Wilde AD, Snyder DJ, Putnam NE, Valentino MD, Hammer ND, Lonergan ZR, Hinger SA, Aysanoa EE, Blanchard C, Dunman PM, Wasserman GA, Chen J, Shopsin B, Gilmore MS, Skaar EP, Cassat JE. 2015. Bacterial hypoxic responses revealed as critical determinants of the host-pathogen outcome by TnSeq analysis of *Staphylococcus aureus* Invasive Infection. PLoS Pathog 11:e1005341.

67. Duthie ES, Lorenz LL. 1952. Staphylococcal coagulase; mode of action and antigenicity. J Gen Microbiol 6:95–107.

68. Benson MA, Lilo S, Wasserman GA, Thoendel M, Smith A, Horswill AR, Fraser J, Novick RP, Shopsin B, Torres VJ. 2011. *Staphylococcus aureus* regulates the expression and production of the staphylococcal superantigen-like secreted proteins in a Rot-dependent manner. Mol Microbiol 81:659–675.

69. Benson MA, Lilo S, Nygaard T, Voyich JM, Torres VJ. 2012. Rot and SaeRS cooperate to activate expression of the staphylococcal superantigen-like exoproteins. J Bacteriol 194:4355–4365.

70. Ji G, Beavis R, Novick RP. 1997. Bacterial interference caused by autoinducing peptide variants. Science 276:2027–2030.

71. Baba T, Takeuchi F, Kuroda M, Yuzawa H, Aoki K, Oguchi A, Nagai Y, Iwama N, Asano K, Naimi T, Kuroda H, Cui L, Yamamoto K, Hiramatsu K. 2002. Genome and virulence determinants of high virulence community-acquired MRSA. Lancet 359:1819–1827.

72. Kehl-Fie TE, Chitayat S, Hood MI, Damo S, Restrepo N, Garcia C, Munro KA, Chazin WJ, Skaar EP. 2011. Nutrient metal sequestration by calprotectin inhibits bacterial superoxide defense, enhancing neutrophil killing of *Staphylococcus aureus*. Cell host & microbe 10:158–164.

73. Lauderdale KJ, Boles BR, Cheung AL, Horswill AR. 2009. Interconnections between Sigma B, *agr*, and proteolytic activity in *Staphylococcus aureus* biofilm maturation. Infect Immun 77:1623–1635.

74. Luong TT, Dunman PM, Murphy E, Projan SJ, Lee CY. 2006. Transcription profiling of the *mgrA* regulon in *Staphylococcus aureus*. J Bacteriol 188:1899–1910.

75. Mesak LR, Yim G, Davies J. 2009. Improved lux reporters for use in *Staphylococcus aureus*. Plasmid 61:182–187.

76. Figueroa M, Jarmusch AK, Raja HA, El-Elimat T, Kavanaugh JS, Horswill AR, Cooks RG, Cech NB, Oberlies NH. 2014. Polyhydroxyanthraquinones as quorum sensing inhibitors from the guttates of *Penicillium restrictum* and their analysis by desorption electrospray ionization mass spectrometry. J Nat Prod 77:1351–1358.

77. Kinkel TL, Roux CM, Dunman PM, Fang FC. 2013. The *Staphylococcus aureus* SrrAB two-component system promotes resistance to nitrosative stress and hypoxia. mBio 4:e00696–00613.

78. Chen PR, Nishida S, Poor CB, Cheng A, Bae T, Kuechenmeister L, Dunman PM, Missiakas D, He C. 2009. A new oxidative sensing and regulation pathway mediated by the MgrA homologue SarZ in *Staphylococcus aureus*. Mol Microbiol 71:198–211.

79. Dmitriev A, Chen X, Paluscio E, Stephens AC, Banerjee SK, Vitko NP, Richardson AR. 2021. The intersection of the *Staphylococcus aureus* Rex and SrrAB regulons: an example of metabolic evolution that maximizes resistance to immune radicals. mBio 12:e0218821.

80. Brignoli T, Manetti AGO, Rosini R, Haag AF, Scarlato V, Bagnoli F, Delany I. 2019. Absence of protein A expression is associated with higher capsule production in Staphylococcal isolates. Front Microbiol 10:863.

81. Mashruwala AA, Boyd JM. 2017. The *Staphylococcus aureus* SrrAB regulatory system modulates hydrogen peroxide resistance factors, which imparts protection to aconitase during aerobic growth. PLoS One 12:e0170283.

82 Dyzenhaus S, Sullivan MJ, Alburquerque B, Boff D, van de Guchte A, Chung M, Fulmer Y, Copin R, Ilmain JK, O’Keefe A, Altman DR, Stubbe FX, Podkowik M, Dupper AC, Shopsin B, van Bakel H, Torres VJ. 2022. MRSA lineage USA300 isolated from bloodstream infections exhibit altered virulence regulation. Cell host & microbe.

